# Computational investigation of BMAA and its carbamate adducts as potential GluR2 modulators

**DOI:** 10.1101/2024.01.12.575281

**Authors:** Isidora Diakogiannaki, Michail Papadourakis, Vasileia Spyridaki, Zoe Cournia, Andreas Koutselos

## Abstract

Beta-N-methylamino-L-alanine (BMAA) is a potential neurotoxic non-protein amino acid, which can reach the human body through the food chain. When BMAA interacts with bicarbonate in the human body, carbamate adducts are produced, which share high structural similarity with the neurotransmitter glutamate. It is believed that BMAA and its L-carbamate adducts bind in the glutamate binding site of ionotropic glutamate receptor 2 (GluR2). Chronic exposure to BMAA and its adducts could cause neurological illness such as neurodegenerative diseases. However, the mechanism of BMAA action and its carbamate adducts bound to GluR2 has not been yet elucidated. Here, we investigate the binding modes and the affinity of BMAA and its carbamate adducts to GluR2 in comparison to the natural agonist, glutamate, to understand whether these can act as GluR2 modulators. Initially, we perform Molecular Dynamics (MD) simulations of BMAA and its carbamate adducts bound to GluR2 to examine the stability of the ligands in the S1/S2 ligand-binding core of the receptor. In addition, we utilize alchemical free energy calculations to compute the difference in the free energy of binding of the beta-carbamate adduct of BMAA to GluR2 compared to that of glutamate. Our findings indicate that carbamate adducts of BMAA and glutamate remain stable in the binding site of the GluR2 compared to BMAA. Additionally, alchemical free energy results reveal that glutamate and the beta-carbamate adduct of BMAA have comparable binding affinity to the GluR2. These results provide a rationale that BMAA carbamate adducts may be in fact the modulators of GluR2 and not BMAA itself.

## INTRODUCTION

Beta-N-methylamino-L-alanine (BMAA) is a natural, neurotoxic non-protein amino acid that is produced by a range of ecologically diverse phytoplankton groups such as cyanobacteria, diatoms and dinoflagellates.^1^ Such cyanobacteria live in the roots of the cycad tree, whose seeds are a common food source consumed by humans; thus BMAA may end up in the human body through the food chain. In addition, BMAA may reach the human body through the food chain in aquatic systems as BMAA can be transferred from cyanobacteria via zooplankton to organisms at higher trophic levels.^2^ It has been established that BMAA has neurotoxic and neuro-excitatory properties and that is a source of neurodegenerative disorders in humans; specifically, BMAA has been detected in postmortem brain and spinal cord tissues of Amyotrophic Lateral Sclerosis (ALS), Parkinson’s and Alzheimer’s patients.^3^

Neurotoxicity of BMAA is dependent on the presence of bicarbonate, which is produced from the interaction of carbon dioxide with water.^3^ BMAA is non-toxic in a physiological salt solution, but in presence of bicarbonate, which is present in nature and the human body, a carbamylation reaction takes place that produces BMAA carbamate adducts such as alpha-carbamate and beta-carbamate in a ratio of 86:14, respectively.^3^ BMAA and its carbamate adducts share a high structural similarity with the natural agonist glutamate (Figure 1).^4^ Glutamate and structurally similar substances activate several glutamate receptors (GluRs), which act as neurotransmitters in the nervous system.^5^ GluRs are divided into two subcategories, the metabotropic receptors (mGluR) and the ionotropic receptors (iGluR) that include kainate, α-amino-3-hydroxy-5-methyl-4-isoxazolepropionic acid (AMPA) and N-methyl-D-aspartate (NMDA).^6,7^ The activation of iGluRs is important in the development and function of the nervous system, while they are also essential in memory and learning. Dysfunction of iGluRs leads to excitotoxic cell death.^6^ It has been therefore hypothesized that the role of BMAA role in the onset and progression of neurodegenerative diseases might be linked to the dysfunction of iGluRs, as BMAA and its carbamate adducts might bind in the binding site of iGluRs, due to their structural resemblance with glutamate.^8,9^

**Figure 1.**
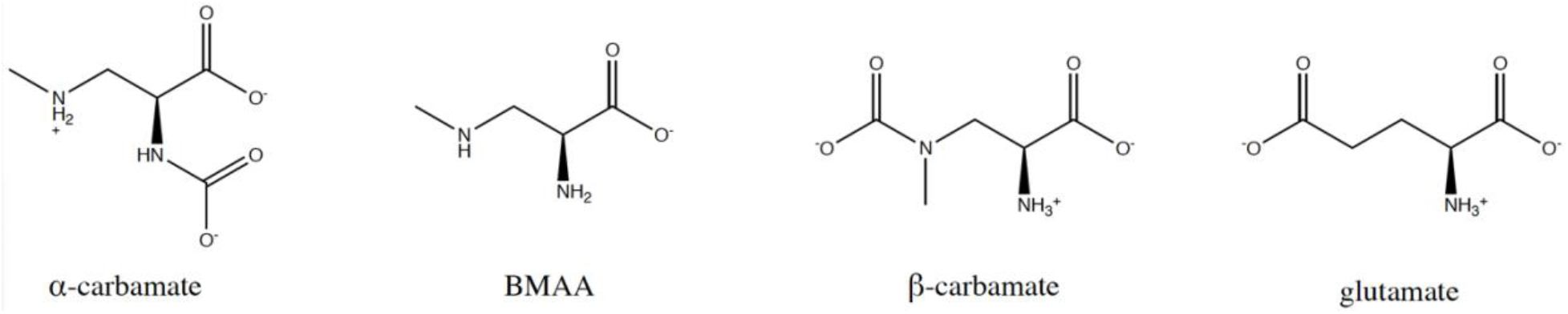
Structures of alpha-carbamate, BMAA, beta-carbamate (L-isomers) and the natural agonist glutamate.

A well-studied subtype of AMPA receptors that may be influenced from the binding of BMAA and its carbamate adducts is the ionotropic glutamate receptor 2 (GluR2).^10^ The binding of glutamate to the GluR2 has been examined extensively both with experimental and computational methods. Crystal structures of glutamate bound to GluR2 (PDB ID: 3DP6^11^, 1FTJ^12^) reveal that the binding pocket for glutamate is situated between the S1 and S2 domains (ligand binding core).^6,12,13^ The IC_50_ for the displacement of ^3^H-AMPA by glutamate has been estimated to be 821 nM in competition binding experiments.^12^ The concentration of the substrate AMPA in these experiments was only 20 nM, which can be considered much lower than K_m_. Because IC_50_ values approximate K_i_ when the concentration used in the assay is much lower than K_m_, we can approximate the glutamate free energy of binding to the GluR2 to be -8.4 kcal/mol (821 nM at 310 K).^12^ The binding free energy of glutamate in the closed conformation of GluR2 was also assessed computationally by Speranskiy and Kurnikova^14^ using the MD/PBSA method and resulting in an estimated value of approximately -13 kcal/mol. Furthermore, Mamonova et al^15^ utilized umbrella sampling and Molecular Dynamics (MD) simulations to describe how the GluR2 ligand binding core interacts with glutamate as well as the conformational changes upon ligand binding. One simplified representation of this process envisioned the protein resembling a clamshell, which encloses glutamate and seals the cleft formed by two lobes connected by a hinge^15^ <u>Taking</u> into account the protein reorganization energy, as determined in this study to be approximately -4 kcal/mol, along with the previously computed binding free energy of glutamate in the closed conformation of GluR2, an estimation of the total binding energy between glutamate and the GluR2 S1S2 ligand binding core was computed to be -9 kcal/mol. In addition, the standard binding free energy of glutamate (along with four other ligands) to the GluR2 has also been calculated using free energy perturbation calculations (FEP) coupled with MD simulations.^16^ The standard binding free energy for glutamate to S1/S2 GluR2 using PDB ID: 1FTJ has been estimated to be - 8.5 ± 1.8 kcal/mol, in agreement with experimental results from competition assays with ^3^H-AMPA (-8.4 kcal/mol, see discussion above).^16^ Finally, atomistic MD simulations have also been employed to investigate the mechanism by which glutamate binds to GluR2.^17^ Two possible pathways with which glutamate traverses during ligand binding have been identified using the string method, and the overall free energy difference between the initial unbound state (protein in the apo state, PDB: 1FTO^12^) and the final bound state in pathways 1 and 2 has been estimated to be -5.8 kcal/mol and -8.8 kcal/mol, respectively. The pathway with the lowest free energy of binding corresponds to the glutamate binding pose with PDB ID: 1FTJ, which is the structure that we also used in our relative binding free energy (RBFE) calculations herein.

While studies suggest that the toxic activity of BMAA is linked to the binding of BMAA and its carbamate adducts to the GluR2 glutamate binding site, to the best of our knowledge, there are no previous computational or experimental studies on the binding of these molecules to GluR2 and their mechanism of binding and energetics have not been yet elucidated^8,9^. Therefore, we set out to investigate the stability and the binding affinity of BMAA and its carbamate adducts (L-isomers) to GluR2 in comparison to the natural agonist, glutamate, to assess whether these molecules could bind on GluR2. If their affinity to GluR2 is similar to or stronger than glutamate affinity, BMAA and its carbamate adducts could potentially provoke dysfunction in neurons, leading to neurodegenerative diseases through binding to the GluR2. Herein, we study the stability of BMAA and its adducts in GluR2 using atomistic MD simulations and RBFE calculations using two force fields to quantify the binding of the beta-carbamate adduct of BMAA with respect to glutamate. Our findings suggest that BMAA carbamate adducts may be in fact the modulators of GluR2 and not BMAA itself.

## METHODS

### Ligand Structural Similarity and Selection of Crystal Structure for the Calculations

To examine the two-dimensional (2D) structural similarity of BMAA and its carbamate adducts with respect to the natural agonist glutamate, the Tanimoto coefficient^17^ (Tc) of each ligand compared to glutamate was calculated using ChemBioServer 2.0^18^ (see the Supporting Information (SI)). In addition, we also computed the three-dimensional (3D) similarities between the compounds using the Shape Screening module (Maestro, Schrödinger, Inc)^19^ and the Pharmacophore type scoring function provided by Phase^20,21^.

The receptor crystal structure should be suitable for atomistic simulations of the protein-ligand complex;^22^ here, the glutamate-GluR2 complex structure was selected based on the following criteria: 1) resolution lower than 3.0 Å and 2) crystallization conditions close to body temperature (T = 310 K) and pH (pH =7.4) (see also the SI). The crystal structure with PDB ID: 1FTJ^12^ meets the above criteria as it includes the S1/S2 ligand binding core of GluR2 in complex with glutamate and it is crystallized at 277 K, pH = 6.5 and resolution = 1.9 Å. Moreover, it has been successfully used in recovering the energetics of glutamate binding using computational work, in excellent agreement with experiments (see the Introduction). Because the crystallized protein with PDB ID 1FTJ is derived from *Rattus norvegicus*, we compared the rat protein sequence with a human protein sequence using BLAST^23^, which showed a 99.5% sequence similarity between the two proteins. The only amino acid change on the S1/S2 core is the G231R mutation that is located at ca. 17 Å from the center of mass of the native agonist, glutamate. We thus concluded that PDB ID 1FTJ^12^ is a suitable starting structure for performing MD/alchemical free energy calculations with glutamate as a ligand. Chain A of the GluR2 ligand binding core was isolated from the full trimer to speed up the simulations (Figure 2). Full details of the model construction and protein modeling are presented in the SI.

**Figure 2.**
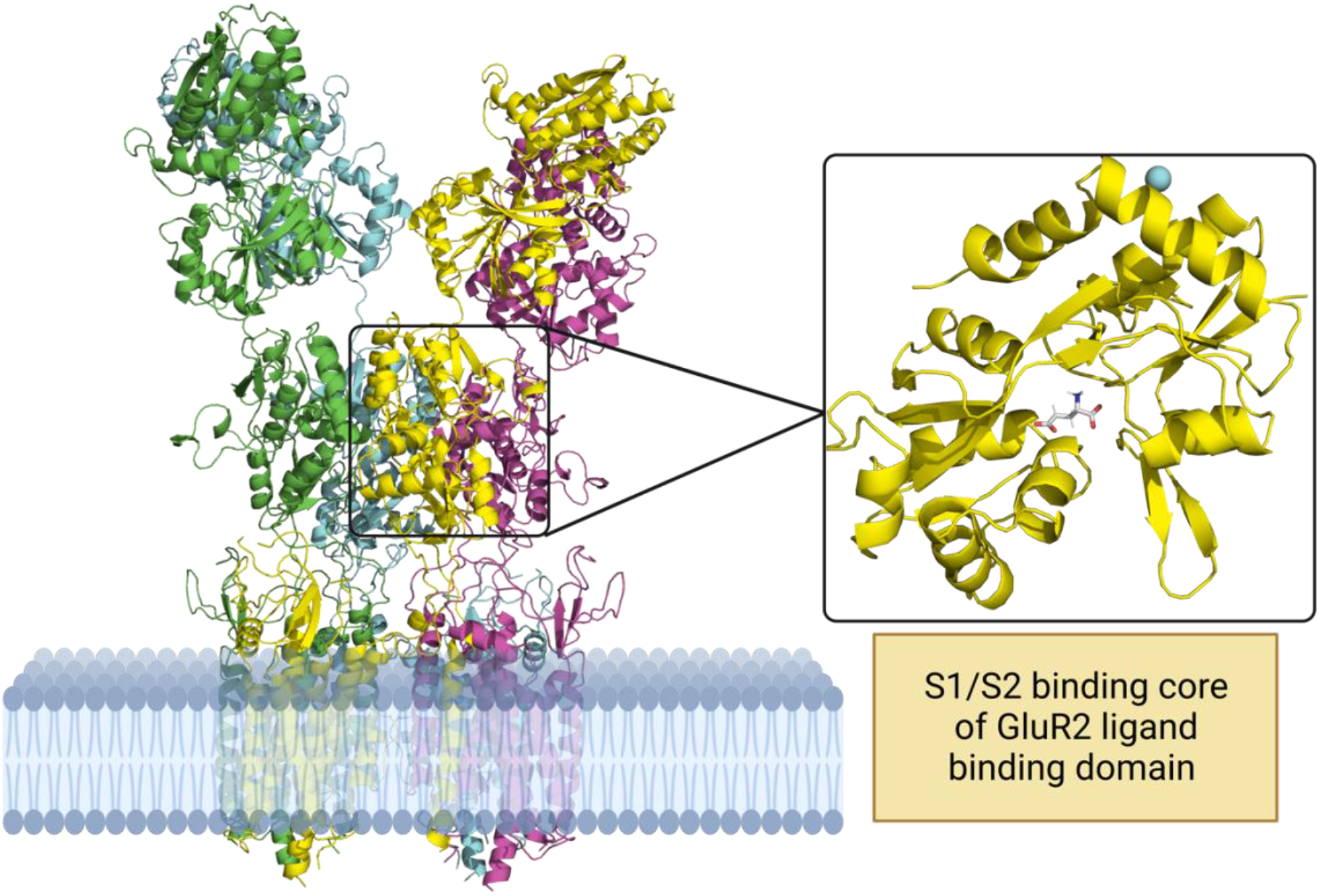
Structure of GluR2 embedded in a phospholipid bilayer. Inset: chain A of the S1/S2 ligand binding core of GluR2 (PDB ID: 1FTJ) is illustrated together with the bound native agonist glutamate shown in stick representation.

### Docking

Before creating starting configurations for BMAA and its carbamate adducts binding on GluR2, we assessed the ability of Glide 6.7^24–26^ to reproduce the glutamate crystal structure pose in the binding pocket of GluR2. Self-docking of glutamate was performed using default parameters of Glide 6.7 (Figure S1). BMAA, alpha-carbamate and beta-carbamate were also docked into GluR2 using default parameters of Glide 6.7 to generate initial positions for the MD simulations. Docking scores were calculated by using the Glide 6.7 SP scoring function.

### MD Simulations with NAMD2.14/OPLS-AA/M/CM1A

Three replicas for each system, namely glutamate/GluR2, BMAA/GluR2, alpha-carbamate/GluR2 and beta-carbamate/GluR2, were simulated for 100 ns using the NAMD 2.14 software. For the replica simulations all conditions were identical except for different initial velocities. For the protein, the OPLS-AA/M^27^ force field was used. Ligand parameters were retrieved from the LigParGen^28^ Web server (OPLS-AA/CM1A^29^ force field) and the TIP3P^30^ potential was used for water. The systems were solvated in a cubic box of 15 Å length in each dimension with 45,891 water molecules. Counter ions were added to neutralize the total charge. Long-range electrostatic interactions were treated using the particle-mesh Ewald (PME) summation method^31^. A time step of 2 fs was used. The temperature was kept constant at 310 K using the Langevin thermostat^32^ with a time constant of 1 ps. The pressure was isotropically maintained at 1 atm using the Nosé-Hoover barostat^33^. The barostat oscillation period was set to 100 fs and the barostat damping timescale was set to 50 fs. The non-bonded potential energy functions (electrostatic and van der Waals) were smoothly truncated between cutoff distances of 10-12 Å.

Prior to MD simulations, all structures were relaxed with 50,000 steps of energy minimization using steepest descent. Heating followed, where the temperature was incrementally increased from 10 to 310 K for 0.6 ns. Then, the systems were equilibrated in the NPT ensemble for 1 ns. Finally, unbiased MD simulations were carried out for 100 ns. To assess the stability of the ligands in the binding cavity of the GluR2, the RMSD of each ligand was calculated with respect to the initial frame using the *RMSD Trajectory tool* of VMD 1.9.3^34^.

### MD Simulations with AMBER20/ff19SB/GAFF2

The same procedure for model construction, equilibration, and MD production protocols was followed using the AMBER20^35^ software with the ff19SB^36^ force field for the protein and GAFF2^37^ parameters that use AM1-BCC^38^ charges for the ligands. The Monte Carlo barostat^39^ of AMBER20 was used and the pressure was set to 1 atm. The number of steps between the performed volume change attempts was equal to 100.

### Relative binding free energy calculations

The relative binding free energy of glutamate bound on GluR2 compared to beta-carbamate was computed with alchemical relative free energy calculations^40–42^. An example of the thermodynamic cycle used for this study is illustrated in Figure 3. For more information on alchemical free energy theory, please refer to the SI or recent reviews.^41^

**Figure 3.**
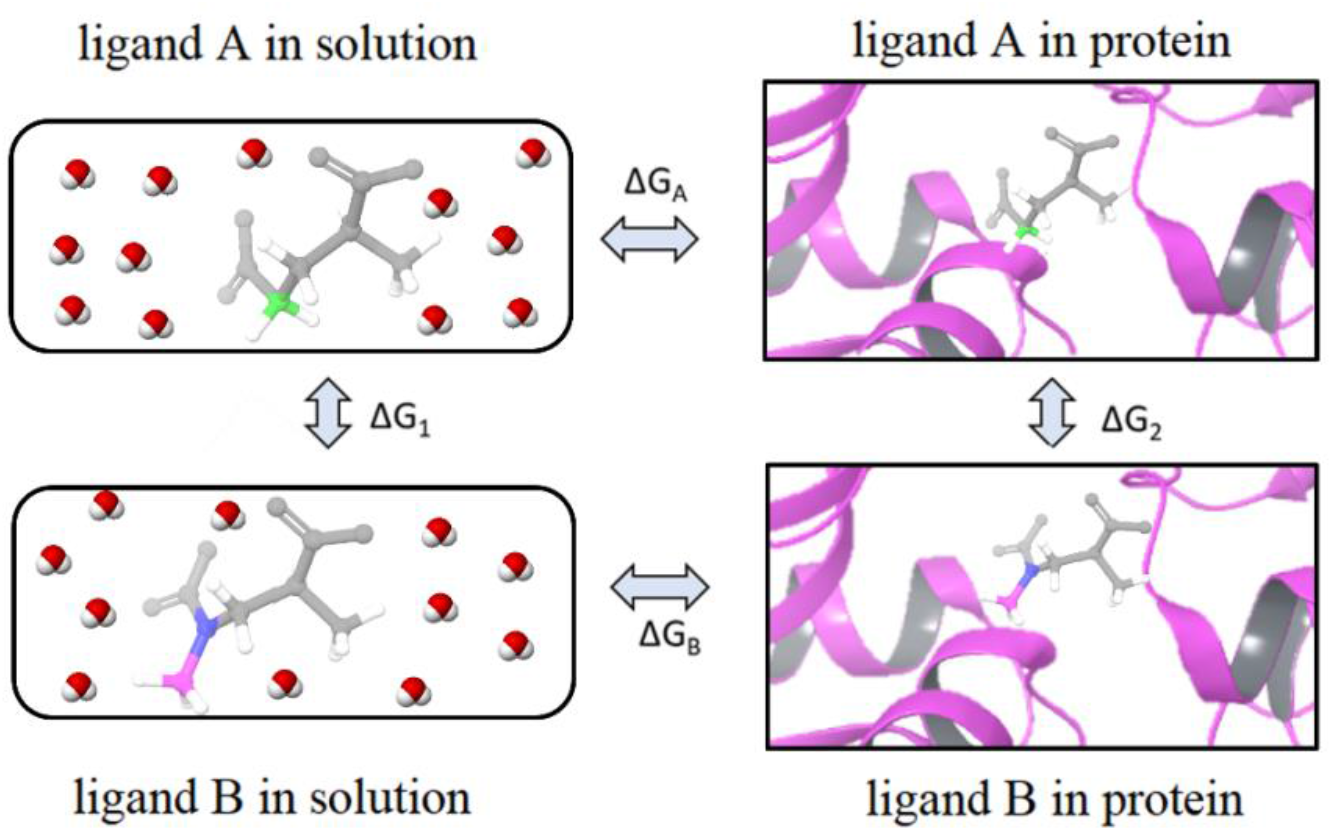
Thermodynamic cycle for the perturbation of ligand A (glutamate) to ligand B (beta-carbamate), in which a methylene bridge colored in green is replaced by a methylamino group colured in blue and pink. All other atoms are colored in gray. The relative binding free energy (ΔΔG_bind_) of beta-carbamate with respect to glutamate can be calculated via two possible paths. Simulating the direct path (horizontal processes, ΔG_B_ - ΔG_A_) is slow to converge due to the large difference between the end states (solution versus protein environments). The alchemical path (vertical processes, ΔG_2_ – ΔG_1_) requires much smaller perturbations to the system and therefore tends to converge much faster. Here, ligand A (glutamate) is perturbed to ligand B (beta-carbamate) in the bound state (ΔG_2_) and the unbound state (ΔG_1_), the difference of which is identical to the direct path (due to the closed thermodynamic cycle). The double arrow for each process illustrates that both forward and backward perturbations are performed.

The perturbation between two ligands, in our case glutamate and beta-carbamate, is typically performed by connecting the two systems using a coupling parameter λ, such that the potential energy function U(λ) interpolates between the initial state (glutamate, λ=0.0) and the final state (beta-carbamate, λ=1.0). To increase the phase space overlap between the two ligands, we introduced two intermediate molecules, named intermediate-1 and intermediate-2 (Figure 4). In intermediate-1 a methylene bridge of glutamate is replaced by an amino group, and in intermediate-2 the nitrogen of the methylamino group of beta-carbamate is replaced by a carbon-hydrogen bond.

**Figure 4.**
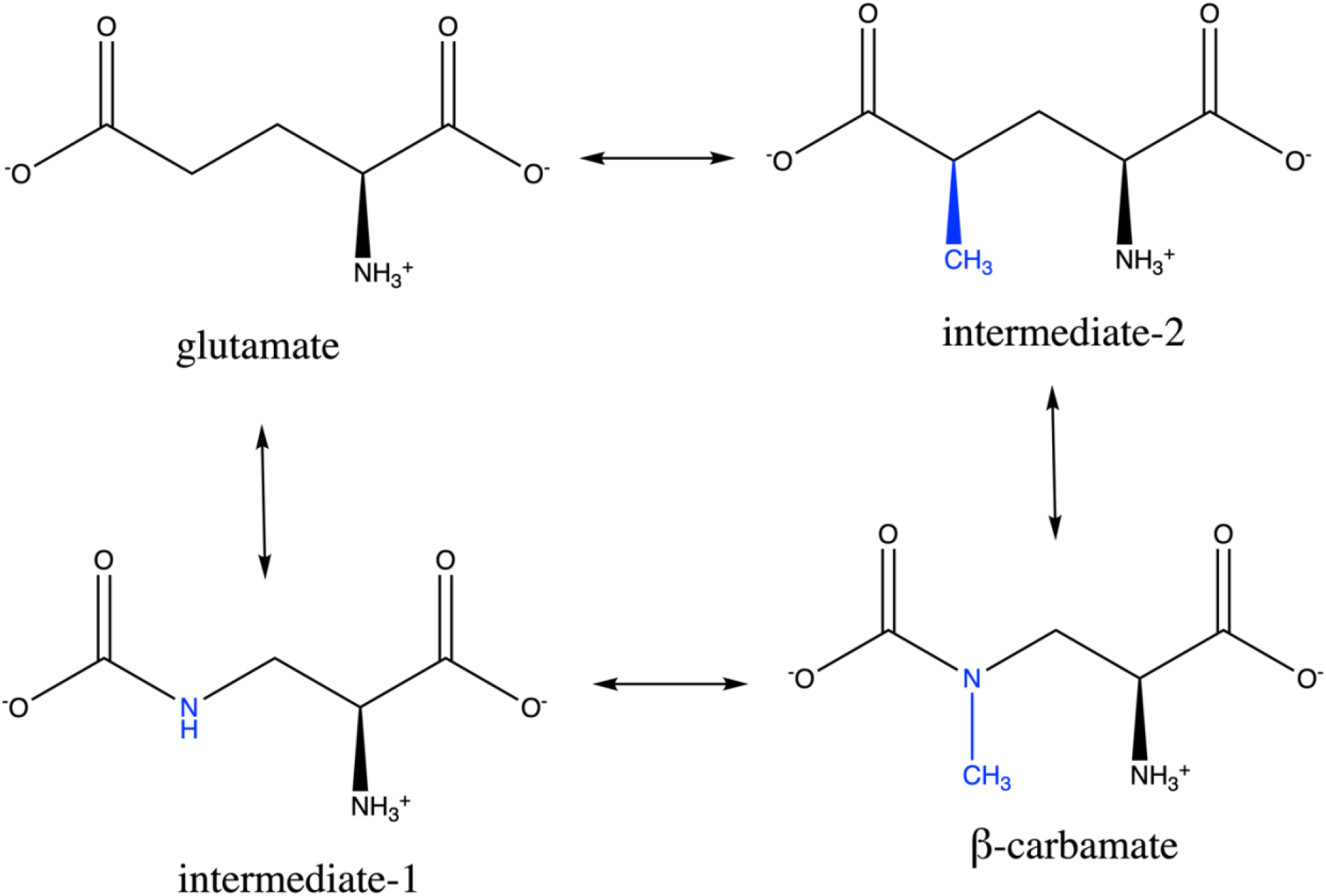
Perturbation network used for the RBFE calculations. Glutamate and beta-carbamate are connected through two intermediate molecules to increase phase space overlap and create a closed cycle. The perturbed atoms with respect to glutamate are coloured blue.

In the generated relative binding free energy (RBFE) network each edge represents both forward and backward perturbations that are performed in both the bound and the unbound state. Moreover, a closed thermodynamic cycle is constructed, where hysteresis can be measured because the summation of the free energy change along each edge of the closed cycle should be zero.

The primary assumption of RBFE calculations is that all ligands in the congeneric series retain the same binding mode. Therefore, via Maestro Schrödinger Suite 2020-3,^19^ beta-carbamate, intermediate-1 and intermediate-2 were overlaid using as a starting structure the crystallographic binding orientation of glutamate into chain A of GluR2 (Figure S2).

### FEP/MD simulations using NAMD 2.14/OPLS-AA

For FEP/MD simulations using NAMD 2.14/OPLS-AA, we generated all input files using the FEPrepare Web server^43^, which automates the setup procedure for relative binding FEP calculations for the dual-topology scheme of NAMD using OPLS-AA force field topology and parameter files. Then, all structures were equilibrated in the *NVT* and *NPT* ensembles for 1 and 2 ns, respectively. Production run simulations were run for 10 ns with NAMD 2.14 in the NPT ensemble per λ window. The number of equidistant λ windows employed for each perturbation was 17 (values for each window ranged between 0.00 and 1.00). A 2 fs timestep was used and a softcore potential was applied to keep pairwise interaction energies finite for all configurations and provide smooth free energy curves for all the simulations^44,45^. A Langevin thermostat and a Nosé-Hoover barostat were applied for the control of the temperature and pressure, respectively. The damping coefficient for Langevin dynamics was set at 1 ps, while the barostat oscillation period was set at 100 fs, and the barostat damping timescale was set to 50 fs. Finally, a 12 Å atom-based cutoff distance for the non-bonded interactions was used and Coulombic interactions were handled using PME.

### TI/HREMD simulations using AMBER20/ff14SB

AMBER20 input files were generated using the ProFESSA workflow,^46^ which automates the setup procedure for the alchemical enhanced sampling (ACES^47^) protocol implemented in the GPU-accelerated AMBER free energy MD engine. In this protocol, a two-state simulation setup in conjunction with Hamiltonian replica-exchange molecular dynamics (HREMD^48^) was employed to increase the conformational sampling of the alchemical free energy calculations.^49^ The equilibration protocol was divided into two phases; the first phase consisted of rigorous equilibration of the initial (λ=0) and end (λ=1) states. First, a minimization with Cartesian restraints relative to the starting structure was implemented to all non-solvent atoms with a force constant of 5 kcal/mol/Å^2^ for 5000 steps followed by 5000 steps of minimization of the full system without any restraint. Then, the same restraints were employed for the system to perform an NPT equilibration for 1.2 ns. Subsequently, the system was heated at a fixed volume at 310 K for 1 ns followed by 1 ns equilibration with the NPT ensemble. Then, annealing was performed, where the system was heated to 600 K for 0.1 ns, simulated at 600 K for 0.3 ns and finally cooled down to 310 K in the last 0.4 ns. At that point, the Cartesian restraints were gradually released during a series of simulations of 0.4 ns with the NPT ensemble. Finally, the system was simulated with no restraints with an NPT ensemble for an additional 0.4 ns.

A second phase of equilibration simulations then followed, where 25 λ states were generated and equilibrated independently. The first half of the λ windows were generated from the equilibrated λ=0 state and the second half of the λ windows were generated from the equilibrated λ=1 phase. The full system was minimized for 5000 steps using the steepest descent algorithm, then heated at fixed volume with 310 K for 1 ns and equilibrated in the NPT ensemble for 4 ns. Finally, production runs were initiated from the final equilibrated structures, and 10 ns MD simulations were performed for each λ window using a 2 fs timestep. A recently developed robust smoothstep softcore potential was applied to avoid the end-point catastrophe, particle collapse and large gradient-jump problems often encountered in alchemical free energy simulations^50^. For temperature control at 310 K, a Langevin thermostat was employed with a friction constant of 2.0 ps^-1^. The pressure was maintained at 1 atm using a Monte Carlo barostat (100 time steps between isotropic box scaling attempts). For the short non-bonded interactions, a 12 Å atom-based cutoff distance was used, while the long-range electrostatics were evaluated with the PME method. The AMBER20/ff14SB/thermodynamic integration (TI) protocol was performed three times, using different initial velocities drawn from the Maxwell-Boltzmann distribution.

### Analysis of RBFE calculations

To estimate the binding free energy differences from the NAMD2.14/OPLS-AA/FEP simulations, the Multistate Bennett Acceptance Ratio (MBAR^51^) was implemented using the ParseFEP plugin^52^ of VMD 1.9.3. Differences in binding free energies from the AMBER20/ff14SB/TI simulations were also estimated with MBAR using Python scripts kindly provided by the Darrin York group. The reported binding free energies from the AMBER20/ff14SB/TI protocol are the mean of the three runs, while statistical uncertainties are calculated as the standard error of the mean.

## RESULTS AND DISCUSSION

### Ligand Structural Similarity and Generation of Initial Poses

To assess the ability of BMAA and its carbamate adducts to act as GluR2 modulators, we first calculated their 2D structural similarity with respect to glutamate, which revealed a high structural similarity of the molecules under study with the natural agonist with Tc=0.5 (see Methods for a definition of the Tc) for alpha- and beta-carbamates and 0.47 for BMAA (Table S1). Moreover, we performed 3D similarity calculations of BMAA and carbamate adducts using glutamate as the query structure for the Shape Screening calculations. The results showed moderate similarity scores of beta-carbamate (0.53), alpha-carbamate (0.44) and BMAA (0.50) highlighting that beta-carbamate has the highest binding pose similarity compared to the natural agonist (Table S2).

To test whether Glide can produce valid starting structures for MD simulations, we self-docked glutamate to the GluR2 crystal structure (PDB ID: 1FTJ) with Glide 6.7^24–26^; the RMSD between the docked and the crystal glutamate poses was 1 Å (Figure S1) indicating that Glide can accurately predict the binding pose of glutamate.

As starting structures for the MD simulations of BMAA and its carbamate adducts, we used the top docked pose from the Glide docking protocol. Because BMAA has a single acidic group compared to glutamate, the acidic group could potentially bind in both directions adopting two flipped binding modes. To investigate the possibility of a flipped BMAA binding pose compared to the top Glide docked pose, we exported the first 13 poses from the Glide docking protocol. From the 13 poses, only the 7th (ΔG = -4.04 kcal/mol) and 8th (ΔG = -3.92 kcal/mol) docking pose bound the acid in the opposite direction, having a ca. 2.39 kcal/mol difference from the top docked pose (ΔG = -6.39 kcal/mol). Therefore, we proceeded with the most energetically favourable docking pose of BMAA and its carbamate adducts to perform the MD simulations.

### MD simulations

We then investigated the binding of BMAA and its carbamate adducts to GluR2 by performing 100 ns equilibrium MD simulations of glutamate, BMAA, alpha-carbamate, and beta-carbamate in complex with the GluR2 using two different protocols: NAMD 2.14^53^ software with the OPLS-AA force-field^27–29^ (NAMD2.14/OPLS-AA) and AMBER20^35^ software using the ff19SB^36^ and GAFF2^37^ force-fields (AMBER20/ff19SB), all in triplicates with identical starting conditions but different initial velocities. Calculation of the trajectory RMSD with respect to the initial frame of the simulation showed that all four molecules remain stable in the binding pocket of the receptor using the NAMD2.14/OPLS-AA protocol (RMSD_glutamate_ = 4.6 ± 1.3 Å, RMSD_BMAA_ = 3.3 ± 0.9 Å, RMSD_alpha-carbamate_ = 2.8 ± 0.8 Å, RMSD_beta-carbamate_ = 3.2 ± 0.8 Å) (Figure S3, Table 1) except for BMAA in replica 1 (RMSD = 28.0 ± 18.1 Å) and beta-carbamate in replica 3 (RMSD = 11.4 ± 10.7 Å) that became solvent exposed at 27 ns and 45 ns, respectively. Using AMBER20/ff19SB, glutamate (RMSD = 1.7 ± 0.7 Å), alpha-carbamate (RMSD = 1.7 ± 0.2 Å), and beta-carbamate (RMSD = 2.5 ± 0.8 Å) remained stable in the binding site of the GluR2 (Figure S4, Table 2) for all simulations. BMAA drifted off the binding site in two out of three replicas, at 74 ns in replica 1 (RMSD = 11.4 ± 15.8 Å) and at 21 ns in replica 3 (RMSD = 30.0 ± 18.4 Å). For the calculation of the average RMSD between the three replicas for both protocols, listed in Table 1 and Table 2, we did not include the RMSD values from the simulations of the dissociated molecules. The average RMSD values obtained from NAMD2.14/OPLS-AA were higher than those observed when using AMBER20/ff19SB with an average difference of ca. 1.50 Å, suggesting that OPLS-AA parameters produce more flexible simulations. Based on our results, we conclude that the substrate glutamate and the ligands alpha-carbamate, and beta-carbamate are stable inside the binding pocket of the GluR2 after 100 ns, while BMAA is less stable and may dissociate from the binding site of the receptor.

**Table 1.** Average values of RMSD and standard deviation of each protein-ligand complex were calculated over the 100 ns MD simulations using NAMD2.14/OPLS-AA.

| Replica ID | RMSD Glutamate (Å) | RMSD BMAA (Å) | RMSD alpha-carbamate (Å) | RMSD beta-carbamate (Å) |
| --- | --- | --- | --- | --- |
| 1 | 5.9 ± 1.4 | 28.0 ± 18.1 | 3.1 ± 1.2 | 2.6 ± 0.3 |
| 2 | 3.3 ± 0.8 | 1.7 ± 0.3 | 2.8 ± 0.4 | 3.8 ± 1.1 |
| 3 | 4.7 ± 1.6 | 4.8 ± 1.3 | 2.4 ± 0.7 | 11.4 ± 10.7 |
| Average RMSD | 4.6 ± 1.3 | 3.3 ± 0.9 | 2.8 ± 0.8 | 3.2 ± 0.8 |

**Table 2.**
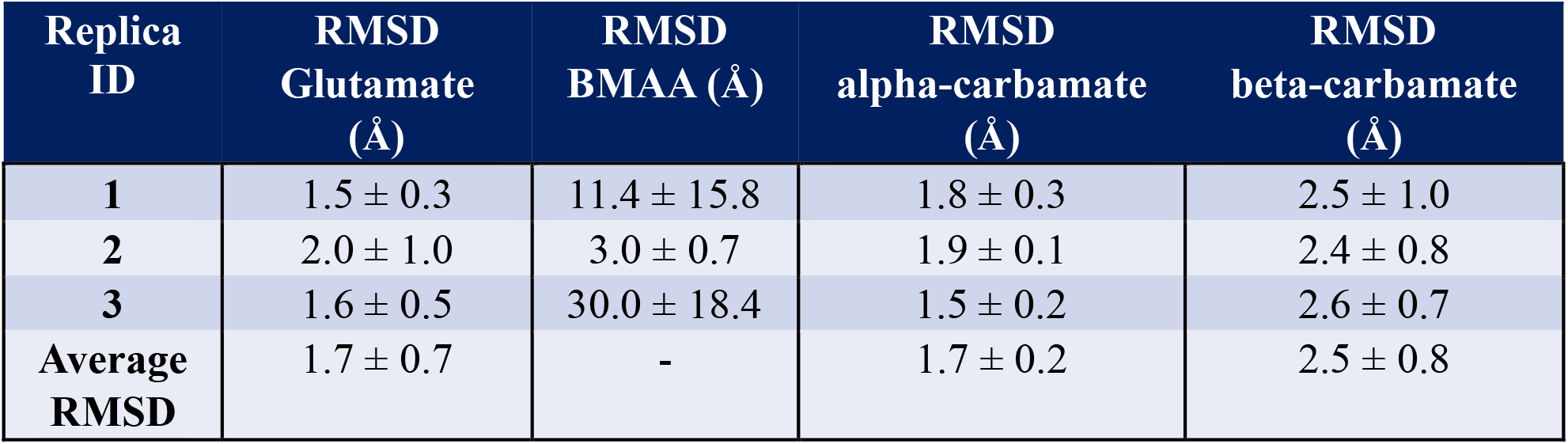
Average values of RMSD and standard deviation of each protein-ligand complex were calculated over the 100 ns MD simulations using AMBER20/ff19SB.

| Replica ID | RMSD Glutamate (Å) | RMSD BMAA (Å) | RMSD alpha-carbamate (Å) | RMSD beta-carbamate (Å) |
| --- | --- | --- | --- | --- |
| 1 | 1.5 ± 0.3 | 11.4 ± 15.8 | 1.8 ± 0.3 | 2.5 ± 1.0 |
| 2 | 2.0 ± 1.0 | 3.0 ± 0.7 | 1.9 ± 0.1 | 2.4 ± 0.8 |
| 3 | 1.6 ± 0.5 | 30.0 ± 18.4 | 1.5 ± 0.2 | 2.6 ± 0.7 |
| Average RMSD | 1.7 ± 0.7 | - | 1.7 ± 0.2 | 2.5 ± 0.8 |

To investigate the structural basis of glutamate and beta-carbamate binding toGluR2 (beta-carbamate is the most structurally similar to glutamate Tc = 0.50 for 2D similarity and Tc = 0.53 for 3D similarity), we used the MDTraj^54^ python package and calculated the pairwise RMSDs between the trajectory conformations to find the first cluster representative from each trajectory. The most important interactions between glutamate and GluR2, which are present in at least three replicas with both simulation protocols, are with amino acids Pro89, Thr91, Arg96, Ser142 and Glu193 (Figure 5A). Dipole-ion interactions form between the carboxylate of glutamate, the Ser142 amide group and the Ser142 side chain hydroxyl group. Glutamate also forms ion-dipole interactions with its amino group and Pro89, Thr91 and Glu193. Salt bridges are present between the glutamate charged amino group and the side chain of Glu192 as well as between the positively charged nitrogen of the guanidinium group of Arg96 and the negatively charged backbone carboxyl oxygen of glutamate. Beta-carbamate’s most important interactions with GluR2 also form with residues Pro89, Thr91, Arg96, Ser142 and Glu193 (Figure 5B). Replica 3 from NAMD2.14/OPLS-AA protocol was not taken into consideration, as beta-carbamate dissociates from the binding site of the receptor. Beta-carbamate forms ion-dipole interactions with Pro89, Thr91, Ser142 and Glu193. A salt bridge forms between the positively charged nitrogen of the guanidium group of Arg96 and the backbone negatively charged carboxylate of beta-carbamate. Ion-dipole interactions of glutamate with Thr91 are present across all replicas for both protocols and the corresponding beta-carbamate/Thr91 interactions, apart from the NAMD2.14/OPLS-AA replica2 simulation. The salt-bridge interaction of beta-carbamate with Arg96 is conserved in all MD simulations, whereas the glutamate/Arg96 interaction is not formed in NAMD2.14/OPLS-AA replica2 simulation. Therefore, a comparison of the structural basis of binding to GluR2 between glutamate and beta-carbamate highlights the high similarity of the formed interactions between the two ligands.

**Figure 5.**
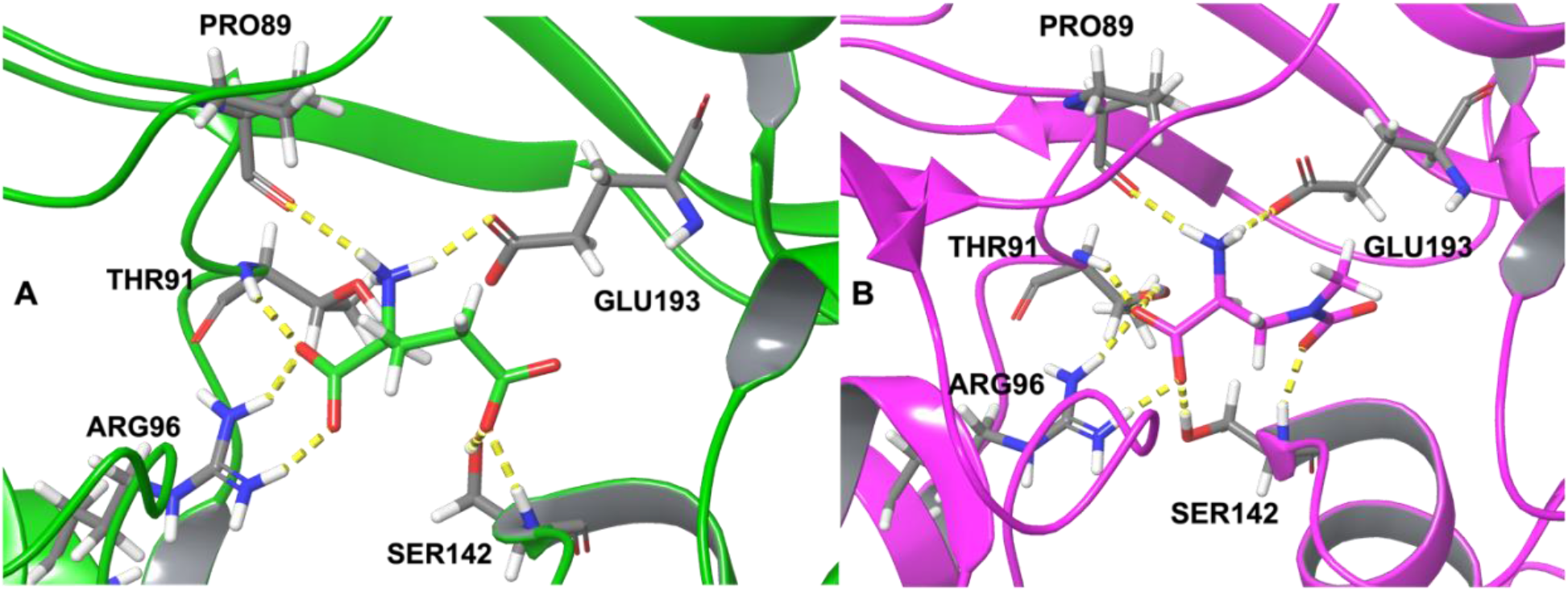
Interactions between A) glutamate and GluR2 and B) beta-carbamate and GluR2.

### RBFE calculations

The relative binding free energy of beta-carbamate with respect to the natural agonist glutamate in GluR2 was determined by alchemical free energy calculations (Figure 3). We examined beta-carbamate because it is the most structurally similar to glutamate (Tc = 0.50 for 2D similarity and Tc = 0.53 for 3D similarity) and remained bound in the binding site of GluR2 during the MD simulations. Prior to the RBFE calculations, we examined whether the beta-carbamate could adopt multiple binding modes within the pocket of GluR2 using Glide docking calculations. We exported 16 docking poses and observed that the 3rd (ΔG = -7.35 kcal/mol), 8th (ΔG = -5.75 kcal/mol), and 10th (ΔG = -5.20 kcal/mol) docking poses flip with respect to the glutamate PDB ID 1FTJ pose. The flipped binding pose with the most favorable free energy had a 1.27 kcal/mol difference from the dominant binding pose (ΔG = -8.62 kcal/mol). Therefore, the flipping of beta-carbamate during the course of the RBFE calculations should be carefully considered. We thus performed two different protocols: FEP coupled with MD simulations using the NAMD 2.14^53,55^ software and the OPLS-AA^27–29^ force-field (NAMD2.14/OPLS-AA/FEP) and Thermodynamic Integration (TI) coupled with MD simulation using the AMBER20^35,56^ software, the ff14SB^57^ force-field for the protein, and the GAFF2^37^ force-field for the ligands (AMBER20/ff14SB/TI). The latter protocol uses HREMD that accounts for flipping of the molecules during the calculations. The ACES protocol implemented in the GPU-accelerated AMBER free energy MD engine has been successfully tested in complex protein-ligand systems that involve ring flips (CDK2) and concerted ligand and protein side-chain conformational changes (T4-lysozyme).^47^ Thus, the AMBER20/ff14SB/TI method may account for a potential flipped binding mode of the beta-carbamate adduct. In addition, AMBER TI RBFE calculations have also been successfully performed in charged ligand perturbations. In the framework of the 2018 Drug Design Data Resource (D3R) grand challenge 4, all the predicted relative binding free energies of the Cathepsin S dataset were within 2 kcal/mol error compared to the experimental results.^58^ The authors concluded that the high accuracy of the predictions resulted from the force field parameters (ff14SB and GAFF); these are also employed in our AMBER20/ff14SB/TI protocol. In addition, RBFE calculations using the same AMBER force fields and TI calculations were also performed in the PTP1B dataset and showed an overall Mean Unsigned Error (MUE) of 1.6 kcal/mol.^59^ This is of critical importance as the binding energetics for glutamate as well as carbamate adducts appear to be primarily driven by electrostatics and atomistic force fields tend to overly reward charge interactions.^60^ Our AMBER20/ff14SB/TI protocol was repeated two more times to ensure reproducibility of the results.

We initially introduced one intermediate molecule (“intermediate 1”) to connect glutamate and the beta-carbamate adduct of BMAA to increase the phase space overlap so as to obtain a more accurate estimate of the relative binding free energy and to close the thermodynamic cycle and measure the hysteresis of the calculation, thus monitoring the convergence of the RBFE calculation (Figure S5). In a converged alchemical free energy calculation, the sum of the free energies in the closed cycle should be zero or close to zero. However, simulations with NAMD2.14/OPLS-AA/FEP resulted in a cycle closure error of -2.11 ± 0.27 kcal/mol. The phase space overlaps between the λ states are listed in Table S3. To overcome this issue, we introduced additional λ windows for each perturbation, where the convergence between neighbouring λ windows was clearly low (overlap < 40%, see SI). The addition of λ windows increased the average phase space overlap of the perturbation glutamate -> intermediate1 by 27.1% (from 33% in the λ windows with overlap < 40% it increased to 59.7%), for intermediate1 -> beta carbamate the overlap increased 29.5% (from 30.30% in the λ windows with overlap < 40% it increased to 59.9%) and for the beta carbamate -> glutamate perturbation it increased 37.2% (from 21.3% in the λ windows with overlap < 40% it increased to 64.41%) (see also the SI for the differences in each perturbation). However, even if the phase space overlap between neighboring windows increased, the cycle closure error changed by only 0.1 kcal/mol. To reduce the cycle closure error and to increase the phase space overlap between the two ligands, we added another intermediate molecule, “intermediate-2” (Figure 4). Although we did not gain a clear benefit in increasing the phase space overlap from adding the second intermediate molecule (the phase space overlap improved in some neighbouring λ windows but deteriorated in others, see SI “Results” section for details), the cycle closure error improved significantly to 1.10 ± 0.09 kcal/mol, compared to the initial three-state RBFE calculations (-2.11 ± 0.27 kcal/mol). Therefore, adding the second intermediate molecule was needed to increase the convergence of the calculation. We thus employed the same scheme in the AMBER20/ff14SB/TI protocol, resulting in a cycle closure error equal to -0.57 ± 0.41 kcal/mol.

To assess the free energy convergence of our calculations, we measured the degree of overlap between the probability distributions characterizing the equilibrium ensembles for the forward and the backward calculations. The phase space overlap between the λ states ranged between 17.7% - 90.9 using NAMD2.14/OPLS-AA/FEP (Figures S6, S8, S10, S12, S14, S16, S18, S20, Table S4), and between 74.5% and 98.3% using AMBER20/ff14SB/TI, except for the transition from the initial state to the next neighboring state (λ_0->1_), which refers to the intermediate-1→beta-carbamate. In this case, the overlap was 21.3, 21,2, 21,2% for replicas 1,2 an 3, respectively, in the solvent and 22.8, 19.7 and 20.0% for replicas 1, 2 and 3, respectively, in the complex system (Figures S6, S8, S10, S12, S14, S16, S18, S20, Table S4), suggesting that more neighboring λ windows could be added in this transformation.

Free energy (ΔG) versus λ for the NAMD2.14/OPLS-AA/FEP protocol and derivative of the gradient of the potential energy (<∂U/∂λ>) as a function of λ versus λ for the AMBER20/ff14SB/TI protocol were also used to determine regions of high curvature across the intermediate λ windows (Figures S7, S9, S11, S13, S15, S17, S19, S21). These regions of phase space serve as a helpful diagnostic as they denote that more sampling or more dense lambda spacing is required to reduce the error of the free energy estimate. In our simulations, we observed low curvature sets of ΔG or <∂U/∂λ> sets, with the possible exception of the λ_0->1_ transition belonging to the intermediate-1→beta-carbamate for both solvent and complex phases. Thus, a good practice would be to ensure a denser λ spacing in this area.

Overall, our results indicate that the beta-carbamate adduct of BMAA is more potent compared to the natural agonist: the difference in free energy of binding (ΔΔG) of beta-carbamate with respect to glutamate is equal to -1.61 ± 0.33 kcal/mol using NAMD2.14/OPLS-AA/FEP protocol and - 2.77 ± 0.12 kcal/mol using AMBER20/ff14SB/TI protocol (Table 3). Therefore, beta-carbamate could potentially bind to GluR2 more strongly than glutamate, potentially leading to dysfunction in neurons and to neurodegenerative diseases. Finally, both protocols showed comparable binding affinities for each perturbation. The bigger differences in binding free energies observed for glutamate -> intermediate 1 (1.34 kcal/mol) and beta-carbamate -> intermediate-2 (1.39 kcal/mol) perturbations can be attributed to the error of the method, which has been determined to be between 1.5-2 kcal/mol, or from the different settings (e.g., force-fields, sampling technique) applied in the two protocols.^41^

**Table 3.**
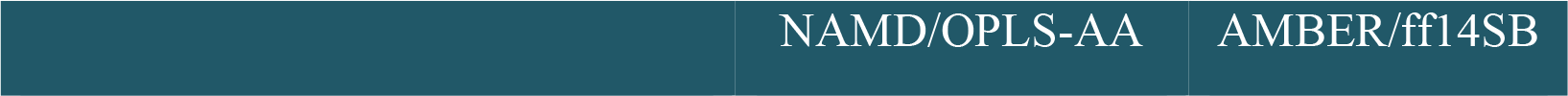
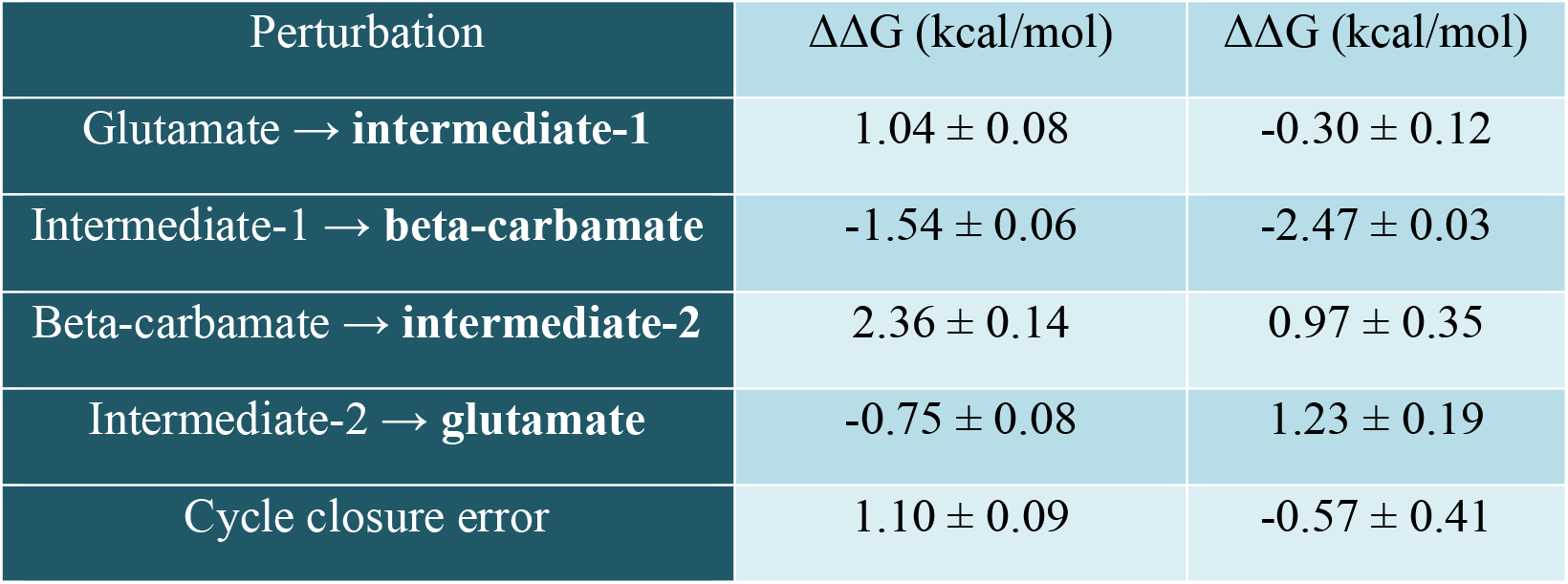
Binding free energy differences for each perturbation and the cycle closure error using NAMD2.14/OPLS-AA and AMBER20/ff14SB.

## CONCLUSIONS

BMAA has been linked to multiple neurodegenerative diseases, but its disease-promoting mechanism remains unknown. BMAA exhibits neurotoxic activity only in the presence of HCO_3_^-^ ions, which leads to the formation of carbamates that share high structural similarity with the glutamic acid. Thus, it has been hypothesized that BMAA and/or its carbamate adducts contribute to neurodegenerative diseases by binding to the glutamate binding site of GluR2.^3^ Although here we do not attempt to describe the disease-promoting mechanism of BMAA, we provide insights into whether BMAA and/or its carbamate adducts bind on GluR2, which could potentially modulate GluR2 at its glutamate binding site and possibly contribute to neurodegenerative diseases.

While the differences between BMAA and its carbamate adducts may look structurally trivial, it is known in the medicinal chemistry literature that small changes in ligand structure may lead to large changes in ligand-protein affinity. Medicinal chemists have long been familiar with the “magic methyl” effect, which is the dramatic change in affinity that may be observed by adding a single methyl group.^61^ Additions of fluorine, nitrogen and other atoms may give rise to the same effect. Because the small changes between glutamate, BMAA and its carbamate adducts cannot be quantified in terms of their affinity for the GluR2 by simply eyeballing the structure, the application of robust and accurate methodologies are needed to calculate the affinity of glutamate compared to its adducts on GluR2, in addition to experimental results to determine these effects.

To investigate whether BMAA and/or its adducts bind on GluR2, we first performed equilibrium MD simulations to monitor the stability of glutamate, BMAA and two carbamate adducts on the GluR2 in atomic-level detail using two different protocols, NAMD2.14/OPLS-AA and AMBER20/ff19SB. Our findings show that the substrate glutamate and the ligands alpha-carbamate and beta-carbamate remain stable in the binding pocket of the GluR2 for 100 ns, while BMAA is less stable and dissociates from the receptor in 50% of the simulations. This observation is in agreement with experimental assays showing that BMAA was non-toxic in the absence of bicarbonate.^3^ Moreover, glutamate and beta-carbamate share similar interactions in the binding pocket of the GluR2, possibly due to their high structural similarity.

Next, to compute the difference in the free energy of binding between the beta-carbamate adduct of BMAA and glutamate to the GluR2, we performed alchemical free energy calculations with two different protocols, NAMD2.14/OPLS-AA/FEP and AMBER20/ff14SB/TI. We did not investigate the binding of BMAA to GluR2 using RBFE calculations because the total charge of these molecules within the perturbation would not remain conserved. Changing the total system charge could yield accurate results,^62^ but there are limited successful examples of charge changes during RBFE calculations in the literature.^41^

Results from the RBFE calculations between the beta-carbamate adduct of BMAA and glutamate binding to GluR2 reveal that using two intermediate molecules instead of one, which was the original setup, significantly improved the closed cycle error of the simulations. The phase space overlap between the different λ states ranged between 17.7% and 90.5% with NAMD2.14/OPLS-AA/FEP and between 74.5% and 98.3% using AMBER20/ff14SB/TI, indicating that the latter protocol was more efficient for this study. In both cases, the difference in the free energy of binding between glutamate and beta-carbamate was negative (-1.61 ± 0.33 kcal/mol for the NAMD2.14/OPLS-AA/FEP protocol and -2.77 ± 0.12 kcal/mol for the AMBER20/ff14SB/TI protocol). Although these values are close to the error of the method, they do suggest that beta-carbamate has a comparable, if not stronger, binding affinity to GluR2 compared toglutamate. Therefore, based on relative binding free energy calculations and studying the interactions between GluR2 and BMAA and its adducts, we provide a rationale that BMAA carbamate adducts may be in fact the modulators of GluR2 and not BMAA itself.

This study also employs two different MD packages and force-fields to ensure the consistency of the results across the codes. Both MD protocols were able to predict the stability of glutamate, alpha-carbamate, and beta-carbamate inside the binding pocket of the GluR2 compared to BMAA. The average RMSD values obtained from NAMD2.14/OPLS-AA were ca. 1.50 Å higher than those observed when using the AMBER20/ff19SB protocol. In addition, the free energy results obtained from the two protocols were also comparable within the limits of the error of the method (1.16 kcal/mol). However, the AMBER20/ff14SB/TI was more efficient for this study based on the phase space overlap between the λ neighbours compared to the NAMD2.14/OPLS-AA/FEP protocol. In addition, the ProFESSA workflow made the AMBER/TI easier to use compared to the NAMD/FEP setup.

Although alchemical free energy calculations are now routinely used in the pharmaceutical industry in lead optimization cycles, they are still not streamlined with non-commercial software. In this paper, we report issues that we faced during the calculations concerning convergence issues and how we solved them (introduction of two intermediate molecules and monitoring the hysteresis of the calculation). We conducted a technically sound study using two different alchemical free energy protocols and given the high interest of the pharmaceutical industry in alchemical free energy calculations, we strongly believe that such studies are important to be demonstrated in the current literature. Especially for the AMBER20/ff14SB/TI protocol, we employed a state-of-the-art methodology that was published recently ^46,47,49^, we tested it and provided a proof of concept for its use. Finally, we provide the full dataset for this protocol, as well as for all the other MD and RBFE protocols that we applied.

Overall, both MD protocols highlight the stability of the beta-carbamate/GluR2 complex, while both RBFE protocols predict a comparable binding affinity of the beta-carbamate and the natural agonist glutamate. Therefore, the beta-carbamate adduct of BMAA could potentially bind equipotently or more strongly to the GluR2 compared to glutamate and may provoke dysfunction in neurons, leading to neurodegenerative diseases; this hypothesis should be further examined through experimental assays.

## SUPPORTING INFORMATION

The Supporting Information is available free of charge.

Additional information and supporting figures and tables describing the ligands’ similarity with glutamate, model construction, relative binding free energy calculations’ theory and results concerning self-docking, and alchemical free energy calculations starting structures and results. (PDF)

## AUTHOR INFORMATION

### Authors

**Isidora I. Diakogiannaki -** Biomedical Research Foundation, Academy of Athens, 4 Soranou Ephessiou, 11527 Athens, Greece, Department of Chemistry, Physical Chemistry Laboratory, National and Kapodistrian University of Athens, Panepistimiopolis, 15771 Athens, Greece

**Michail Papadourakis** *- Biomedical Research Foundation, Academy of Athens, 4 Soranou Ephessiou, 11527 Athens, Greece*

**Vasileia Spyridaki** *- Biomedical Research Foundation, Academy of Athens, 4 Soranou Ephessiou, 11527 Athens, Greece*

### Present Addresses

**Isidora I. Diakogiannaki** – Department of Chemistry, University of Naples Federico II, 49 Via Domenico Montesano, 0810678102 Naples, Italy

### Author Contributions

I.D., M.P. and V.S. performed computational experiments. I.D., M.P., Z.C. and A.K. wrote and revised the manuscript. Z.C. and A.K. conceived the study and supervised the research. All authors have given approval for the final version of the manuscript.

### Notes

The authors declare no competing financial interest.

#### Data and Software Availability

The NAMD 2.14 software package is available at https://www.ks.uiuc.edu/Research/namd/ and the AMBER20 software package is available at http://ambermd.org/. FEPrepare webserver is freely accessible at https://feprepare.vi-seem.eu/ and the LigParGen webserver is freely accessible at http://zarbi.chem.yale.edu/ligpargen/. The input/output files generated during this study are available at https://doi.org/10.5281/zenodo.7730768.

## ACKNOWLEDGMENT

We would like to thank Dr. Anastasia Hiskia for useful discussions about the possible implication of BMAA in GluR2 biology. Thanks are also extended to the Darrin York group and especially Abir Ganguly, Shi Zhang and Darrin York, for useful discussions and for providing the ProFESSA workflow, analysis scripts and the AMBER20 software license for AMBER/ff14SB/TI and AMBER/ff19SB protocols. Finally, we want to thank the reviewers who helped in improving the manuscript.

