## Supplementary Information for "Computational investigation of BMAA and its carbamate adducts as potential GluR2 modulators"

### METHODS

#### Selection of target receptor

From a total of 100 PDB structures available for GluR2, we retained structures that did not contain any mutations and that are complexed with the native substrate/agonist glutamate (eight structures). We then applied the following filtering criteria to select the glutamate-GluR2 complex for the atomistic simulations: 1) resolution lower than 3.0 Å, 2) crystallization conditions close to body temperature (T = 310 K) and pH (pH =7.4). The crystal structure with PDB ID: 1FTJ[1] met the above criteria as it includes the S1/S2 ligand binding core of GluR2 in complex with glutamate and it is crystallized at 277 K, pH = 6.5 and resolution = 1.9 Å. Moreover, this structure has been previously employed in previous computational studies (see the main text). Thus, PDB ID: 1FTJ was selected to build the atomistic model for our simulations.

#### Calculation of BMAA analog similarity with glutamate

To calculate whether alpha-, beta-carbamate and BMAA are structurally similar with respect to the native agonist glutamate, we used the Tanimoto coefficient[2] (*Tc*), a similarity metric for comparing chemical structures using chemical fingerprints. A chemical fingerprint is a series of bits that represent the presence or absence of chemical substructures, patterns, physicochemical characteristics, and other properties in a molecule. If the property is present in the molecule, then the fingerprint is 1 (ON) and if the property is absent, then the fingerprint is 0 (OFF). The *Tc* to compare the similarity of two molecules is given by Equation 1:

$$Tc = \frac{N_C}{N_A + N_B - N_C} \tag{1}$$

where $N_c$ is the number of common ON bits (1) in both molecules, $N_A$ is the number of ON bits in molecule 1 and $N_B$ is the number of ON bits in molecule 2. The *Tc* was calculated using ChemBioServer[3]. The results are presented in Table S1.

**Table S1.** Tanimoto coefficient between glutamate, alpha-carbamate, beta-carbamate and BMAA.

| **Tanimoto coefficient** | | | | |
|---|---|---|---|---|
| Molecule | Glutamate | α-carbamate | β-carbamate | BMAA |
| Glutamate | 1.00 | 0.50 | 0.50 | 0.47 |
| α-carbamate | 0.50 | 1.00 | 0.87 | 0.82 |
| β-carbamate | 0.50 | 0.87 | 1.00 | 0.70 |
| BMAA | 0.47 | 0.82 | 0.70 | 1.00 |

In addition, to compute the 3D similarities among alpha-, beta-carbamate and BMAA with respect to the native agonist glutamate, we used Shape Screening module (Maestro 2020.3, Schrödinger, Inc). 3D similarity screening can be applied to match the overall shape of different compounds[4] by performing initial alignments of atom triplets and then conducting refinement and overlap scoring.[5] To compute the volume overlap scoring between the molecules, we used the Pharmacophore type approach, in which every compound is treated as a collection of pharmacophore sites defined by the Phase[6,7] pharmacophore feature and then calculates the overlapping volumes between same feature sites.[8] A shape similarity score of 0 denotes the maximum dissimilarity between the binding poses of two compounds, while a shape similarity score of 1 means that two compounds have the maximum 3D similarity. The results are presented in Table S2.

**Table S2.** 3D similarity calculations between glutamate, alpha-carbamate, beta-carbamate and BMAA.

| **Type pharmacophore scoring** | | | | |
|---|---|---|---|---|
| Molecule | Glutamate | α-carbamate | β-carbamate | BMAA |
| Glutamate | 1.00 | 0.44 | 0.53 | 0.50 |
| α-carbamate | 0.44 | 1.00 | 0.48 | 0.50 |
| β-carbamate | 0.53 | 0.48 | 1.00 | 0.54 |
| BMAA | 0.50 | 0.50 | 0.54 | 1.00 |

**Glutamate-GluR2 model construction**

The atomistic model of the GluR2 complex was created based on the crystal structure with PDB ID 1FTJ[1], which comprises the GluR2 ligand binding domain (LBD) in complex with glutamate. GluR2 consists of four subunits each one containing the natural agonist glutamate bound to LBD. However, the crystal structure contains only three chains, where chain A includes one zinc ion and chains B and C include two zinc ions each. Zinc ions are not located inside the binding pocket and are at a distance of ca. 17 Å from the center of mass of glutamate. In this study, only chain A was used for the MD simulations/alchemical free energy calculations to reduce the computational effort. Chain A in complex with glutamate was prepared using the Protein Preparation Wizard[9] (Schrödinger 2020-3). Missing residues of chain A were added and refined using Prime[10,11], tautomer/ionization states were assigned and the resulting structure was minimized.

**Relative binding free energy calculations**

The relative binding free energy for the glutamate/beta-carbamate perturbation was estimated using alchemical free energy calculations[12]. The free energy of difference between two end states in both complex and solvent phases could, in theory, be calculated by sampling configurations, computing the energies of each microstate, and evaluating the partition function Z as written in Equation 2:

$$\Delta G^{\circ}_{bind} = -k_B T ln K^{\circ}_B = -k_B T \left( ln \frac{Z_{complex}}{Z_{solvent}} \right) \quad (2)$$

where $k_B$ is the Boltzmann constant, T is the temperature of the system and $K^{\circ}_B$ is the binding constant. The computation of the binding free energy depends on solving the different partition functions of the system, for example the $Z_{complex}$ is given by Equation 3:

$$Z_{complex} = \int \exp\left(-\beta U_{complex}(q)\right) dq \quad (3)$$

where $U_{complex}$ is the potential energy of conformation q of the two end states in the complex phase. However, direct computation of these partition functions is computationally intractable due to the amount of sampling needed to compute absolute free energies. Instead, it is much easier to compute the differences between the binding affinities of two structural similar compounds using a relative free energy calculations formula derived from Equation 2:

$$\Delta\Delta G^{\circ}_{A,B} = \Delta\Delta G^{\circ}_{B,bind} - \Delta\Delta G^{\circ}_{A,bind} = -k_B T \left( ln \frac{Z^B_{complex} Z^A_{solvent}}{Z^A_{complex} Z^B_{solvent}} \right)$$

$$= \Delta G^{\circ}_{A,B,complex} - \Delta G^{\circ}_{A,B,solvent} \quad (4)$$

This approach depends only on the transformation of system A to system B in the complex and solvent phases and thus it resolves errors associated with computing absolute free energies. This transformation is typically performed by connecting the two systems using a coupling parameter $\lambda$, such that the potential energy function $U(\lambda)$ interpolates between the initial state A ($\lambda$=0.0) and the final state B ($\lambda$=1.0).

To obtain accurate relative binding free energy predictions, a sufficient degree of overlap of the phase space between the initial (A) and the final (B) state is required. This is feasible by introducing a series of N alchemical intermediate $\lambda$ windows along the perturbation path. When sufficient overlap is not attained, one can either rerun the calculations using more $\lambda$ windows or insert intermediate compounds to bridge the conformational space between the two states.

During a relative binding free energy calculation involving multiple compounds, one ligand from the congeneric series acts as a reference to which all other molecules are aligned. As a result, the conserved binding mode as well as the improved overlap between the windows are secured. The reference compound is usually selected as the one that is representative of the ligand set and at the same time with the higher certainty of the predicted binding mode. Then, based on the reference ligand, the compound set forms a network, where the "edges" are the transformations performed using alchemical free energy simulations. The relative binding affinities calculated from the network may contain systematic errors due to inability of the force-field to describe the molecular interactions or/and motions or due to unconverged simulations. Here, an estimation of these errors was obtained using the cycle closure error method as the relative free energy should be zero in a closed cycle.

### RESULTS

#### Extra λ windows in perturbation glutamate → intermediate-1

λ=0.31250 to λ=0.37500 was split: λ=0.31250 to λ=0.34000 and λ=0.34000 to λ=0.37500. The convergence increased from 36.51% to 70.70% and 70.26%.
λ=0.81250 to λ=0.87500 was split: λ=0.81250 to λ=0.84000 and λ=0.84000 to λ=0.87500. The convergence increased from 36.74% to 46.17% and 51.50%.
λ=0.87500 to λ=0.93750 was split: λ=0.87500 to λ=0.90000 and λ=0.90000 to λ=0.93750. The convergence increased from 23.64% to 69.92% and 83.80%.
λ=0.93750 to λ=1.00000 was split: λ=0.93750 to λ=0.97000 and λ=0.97000 to λ=1.00000. The convergence increased from 34.96% to 49.50% and 35.85%.

#### Extra λ windows in perturbation intermediate-1 → β-carbamate of BMAA

λ=0.50000 to λ=0.56250 was split: λ=0.50000 to λ=0.53000 and λ=0.53000 to λ=0.56250. The convergence changed from 28.86% to 27.75% and 72.25%.
λ=0.81250 to λ=0.87500 was split: λ=0.81250 to λ=0.84000 and λ=0.84000 to λ=0.87500. The convergence increased from 31.74% to 64.37% and 75.03%

#### Extra λ windows in perturbation β-carbamate of BMAA → intermediate-2

λ=0.62500 to λ=0.68750 was split: λ=0.62500 to λ=0.65000 and λ=0.65000 to λ=0.68750. The convergence increased from 17.65% to 63.15% and 48.95%.
λ=0.75000 to λ=0.81250 was split: λ=0.75000 to λ=0.78000 and λ=0.78000 to λ=0.81250. The convergence improved from 31.74% to 64.37% and 75.03%
λ=0.87500 to λ=0.93750 was split: λ=0.87500 to λ=0.90000 and λ=0.90000 to λ=0.93750. The convergence increased from 26.08% to 43.62% and 75.80%

**Phase space overlap between neighboring λ windows from NAMD/OPLS-AA/FEP and AMBER/ff14SB/TI protocols**

The results obtained from the NAMD/OPLS-AA/FEP and AMBER/ff14SB/TI protocols concerning the phase space overlap between the neighboring λ windows for each perturbation are listed in Tables S3 – S7:

**Table S3.** Phase space overlap between neighboring λ windows of perturbations glutamate→intermediate-1, intermediate-1→beta-carbamate, beta-carbamate→glutamate. Results obtained from the NAMD2.14/OPLS-AA/FEP protocol before adding additional λ windows.

| | **Glutamate→intermediate-1** | | **intermediate-1→beta-carbamate** | |
|---|---|---|---|---|
| | **Solvent** | **Complex** | **Solvent** | **Complex** |
| **λ** | **overlap (%)** | **overlap (%)** | **overlap (%)** | **overlap (%)** |
| **0→1** | 58.49 | 45.17 | 48.06 | 58.05 |
| **1→2** | 58.49 | 45.17 | 48.06 | 58.05 |
| **2→3** | 54.72 | 46.17 | 40.40 | 59.71 |
| **3→4** | 55.94 | 77.25 | 42.73 | 47.39 |
| **4→5** | 58.16 | 50.06 | 50.61 | 56.16 |
| **5→6** | 60.49 | 36.51 | 46.61 | 66.93 |
| **6→7** | 63.60 | 66.26 | 51.61 | 44.51 |
| **7→8** | 66.93 | 57.82 | 51.50 | 39.84 |
| **8→9** | 51.72 | 62.71 | 31.52 | 28.86 |

| | | | | |
|---|---|---|---|---|
| **9→10** | 59.16 | 87.24 | 57.05 | 73.47 |
| **10→11** | 57.05 | 66.15 | 55.27 | 69.92 |
| **11→12** | 57.94 | 45.73 | 53.83 | 67.70 |
| **12→13** | 54.61 | 47.72 | 56.05 | 62.71 |
| **13→14** | 55.72 | 36.74 | 52.50 | 31.74 |
| **14→15** | 53.61 | 23.64 | 47.06 | 40.40 |
| **15→16** | 50.61 | 34.96 | 46.50 | 53.05 |

| | **Beta-carbamate→glutamate** | |
|---|---|---|
| | **Solvent** | **Complex** |
| **λ** | **overlap (%)** | **overlap (%)** |
| **0→1** | 58.27 | 48.39 |
| **1→2** | 58.28 | 48.39 |
| **2→3** | 58.38 | 48.61 |
| **3→4** | 63.60 | 58.16 |
| **4→5** | 58.38 | 61.60 |
| **5→6** | 25.60 | 49.83 |
| **6→7** | 67.48 | 58.49 |
| **7→8** | 63.60 | 63.37 |
| **8→9** | 60.49 | 56.16 |
| **9→10** | 60.93 | 46.28 |
| **10→11** | 58.93 | 58.05 |

| | | |
|---|---|---|
| **11→12** | 59.49 | 61.04 |
| **12→13** | 58.05 | 41.51 |
| **13→14** | 56.83 | 61.04 |
| **14->15** | 52.28 | 53.61 |
| **15->16** | 49.83 | 41.62 |

**Table S4.** Phase space overlap between neighboring λ windows of perturbations glutamate→intermediate-1, intermediate-1→beta-carbamate, beta-carbamate→intermediate-2, and intermediate-2→glutamate. Results obtained from the NAMD2.14/OPLS-AA/FEP protocol.

| | **Glutamate→intermediate-1** | | **intermediate-1→beta-carbamate** | |
|---|---|---|---|---|
| | **Solvent** | **Complex** | **Solvent** | **Complex** |
| **λ** | **overlap (%)** | **overlap (%)** | **overlap (%)** | **overlap (%)** |
| **0→1** | 58.49 | 45.17 | 48.06 | 58.05 |
| **1→2** | 58.49 | 45.17 | 48.06 | 58.05 |
| **2→3** | 54.72 | 46.17 | 40.40 | 59.71 |
| **3→4** | 55.94 | 77.25 | 42.73 | 47.39 |
| **4→5** | 58.16 | 50.06 | 50.61 | 56.16 |
| **5→6** | 60.49 | 36.51 | 46.61 | 66.93 |
| **6→7** | 63.60 | 66.26 | 51.61 | 44.51 |
| **7→8** | 66.93 | 57.82 | 51.50 | 39.84 |

| | | | | |
|---|---|---|---|---|
| **8→9** | 51.72 | 62.71 | 31.52 | 28.86 |
| **9→10** | 59.16 | 87.24 | 57.05 | 73.47 |
| **10→11** | 57.05 | 66.15 | 55.27 | 69.92 |
| **11→12** | 57.94 | 45.73 | 53.83 | 67.70 |
| **12→13** | 54.61 | 47.72 | 56.05 | 62.71 |
| **13→14** | 55.72 | 36.74 | 52.50 | 31.74 |
| **14→15** | 53.61 | 23.64 | 47.06 | 40.40 |
| **15→16** | 50.61 | 34.96 | 46.50 | 53.05 |
| | **Beta-carbamate→intermediate-2** | | **intermediate-2→glutamate** | |
| | **Solvent** | **Complex** | **Solvent** | **Complex** |
| **λ** | **overlap (%)** | **overlap (%)** | **overlap (%)** | **overlap (%)** |
| **0→1** | 63.26 | 45.06 | 90.57 | 76.91 |
| **1→2** | 56.71 | 66.15 | 87.13 | 46.73 |
| **2→3** | 58.60 | 90.90 | 80.69 | 63.26 |
| **3→4** | 57.82 | 86.57 | 87.13 | 82.69 |
| **4→5** | 55.38 | 51.83 | 86.79 | 56.16 |
| **5→6** | 50.39 | 69.70 | 76.69 | 76.14 |
| **6→7** | 57.38 | 65.04 | 77.80 | 81.69 |
| **7→8** | 59.71 | 77.36 | 69.26 | 44.06 |
| **8→9** | 71.59 | 78.14 | 83.13 | 49.28 |
| **9→10** | 58.16 | 60.49 | 78.47 | 38.96 |

| | | | | |
|---|---|---|---|---|
| **10→11** | 69.26 | 17.65 | 83.57 | 44.17 |
| **11→12** | 56.49 | 59.38 | 76.03 | 72.59 |
| **12→13** | 60.71 | 20.09 | 76.58 | 62.60 |
| **13→14** | 54.94 | 64.37 | 71.03 | 64.59 |
| **14->15** | 49.61 | 26.08 | 72.59 | 50.83 |
| **15->16** | 48.50 | 47.39 | 66.93 | 39.07 |

**Table S5.** Phase space overlap between neighboring λ windows of perturbations glutamate→intermediate-1, intermediate-1→beta-carbamate, beta-carbamate→intermediate-2, and intermediate-2→glutamate. Results obtained from replica 1 of the AMBER20/ff14SB/TI protocol.

| | **Glutamate→intermediate-1** | | **intermediate-1→beta-carbamate** | |
|---|---|---|---|---|
| | **Solvent** | **Complex** | **Solvent** | **Complex** |
| **λ** | **overlap (%)** | **overlap (%)** | **overlap (%)** | **overlap (%)** |
| **0→1** | 93.80 | 93.00 | 21.30 | 22.80 |
| **1→2** | 93.50 | 92.60 | 74.50 | 75.10 |
| **2→3** | 93.20 | 94.00 | 83.50 | 83.80 |
| **3→4** | 94.20 | 93.80 | 86.20 | 87.30 |
| **4→5** | 93.80 | 93.90 | 87.60 | 88.50 |
| **5→6** | 93.70 | 94.60 | 88.70 | 89.00 |
| **6→7** | 94.20 | 94.70 | 89.00 | 89.40 |

| | | | | |
|---|---|---|---|---|
| **7→8** | 93.70 | 94.80 | 89.00 | 89.10 |
| **8→9** | 95.10 | 94.90 | 89.10 | 89.40 |
| **9→10** | 94.80 | 94.80 | 88.40 | 88.40 |
| **10→11** | 95.40 | 94.80 | 88.20 | 88.20 |
| **11→12** | 95.00 | 95.80 | 87.80 | 87.20 |
| **12→13** | 96.10 | 95.10 | 87.10 | 87.10 |
| **13→14** | 96.00 | 95.70 | 85.40 | 85.10 |
| **14→15** | 96.70 | 96.50 | 86.20 | 85.90 |
| **15→16** | 96.80 | 96.60 | 84.80 | 84.60 |
| **16→17** | 96.90 | 96.50 | 83.10 | 82.90 |
| **17→18** | 96.90 | 97.10 | 84.40 | 83.80 |
| **18→19** | 96.60 | 96.40 | 82.10 | 82.40 |
| **19→20** | 96.40 | 96.40 | 82.20 | 81.70 |
| **20→21** | 95.50 | 96.30 | 81.60 | 81.30 |
| **21→22** | 95.00 | 95.00 | 80.40 | 80.60 |
| **22→23** | 91.10 | 94.00 | 80.50 | 80.10 |
| **23→24** | 78.00 | 87.90 | 79.90 | 79.90 |
| | **Beta-carbamate→intermediate-2** | | **intermediate-2→glutamate** | |
| | **Solvent** | **Complex** | **Solvent** | **Complex** |
| **λ** | **overlap (%)** | **overlap (%)** | **overlap (%)** | **overlap (%)** |
| **0→1** | 97.80 | 98.30 | 81.30 | 81.30 |

| | | | | |
|---|---|---|---|---|
| **1→2** | 97.80 | 98.00 | 81.60 | 81.60 |
| **2→3** | 97.90 | 97.60 | 81.90 | 81.30 |
| **3→4** | 97.80 | 97.30 | 83.50 | 82.60 |
| **4→5** | 97.80 | 97.20 | 82.00 | 82.70 |
| **5→6** | 97.00 | 97.30 | 85.30 | 84.20 |
| **6→7** | 96.80 | 97.30 | 85.10 | 84.60 |
| **7→8** | 97.10 | 97.00 | 85.30 | 84.70 |
| **8→9** | 96.60 | 97.20 | 87.30 | 86.50 |
| **9→10** | 96.80 | 97.10 | 86.80 | 86.40 |
| **10→11** | 95.70 | 96.90 | 87.40 | 87.60 |
| **11→12** | 95.60 | 96.20 | 89.80 | 88.80 |
| **12→13** | 94.80 | 95.80 | 90.00 | 89.70 |
| **13→14** | 94.70 | 96.20 | 89.00 | 89.20 |
| **14→15** | 94.60 | 96.00 | 90.80 | 91.20 |
| **15→16** | 94.80 | 95.50 | 90.80 | 90.50 |
| **16→17** | 94.60 | 95.90 | 92.60 | 92.60 |
| **17→18** | 94.00 | 95.40 | 91.20 | 91.80 |
| **18→19** | 93.50 | 95.10 | 93.20 | 93.10 |
| **19→20** | 93.20 | 95.10 | 91.00 | 92.00 |
| **20→21** | 93.30 | 94.10 | 92.50 | 93.20 |

| | | | | |
|---|---|---|---|---|
| **21→22** | 92.90 | 93.30 | 91.30 | 92.40 |
| **22→23** | 94.30 | 94.00 | 90.40 | 91.30 |
| **23→24** | 92.50 | 92.20 | 86.20 | 88.20 |

**Table S6.** Phase space overlap between neighboring λ windows of perturbations glutamate→intermediate-1, intermediate-1→beta-carbamate, beta-carbamate→intermediate-2, and intermediate-2→glutamate. Results obtained from replica 2 of the AMBER20/ff14SB/TI protocol.

| | **Glutamate→intermediate-1** | | **intermediate-1→beta-carbamate** | |
|---|---|---|---|---|
| | **Solvent** | **Complex** | **Solvent** | **Complex** |
| **λ** | **overlap (%)** | **overlap (%)** | **overlap (%)** | **overlap (%)** |
| **0→1** | 93.90 | 93.30 | 21.20 | 19.70 |
| **1→2** | 93.30 | 93.40 | 74.70 | 76.20 |
| **2→3** | 94.20 | 93.50 | 83.20 | 83.90 |
| **3→4** | 93.60 | 93.90 | 86.30 | 87.20 |
| **4→5** | 93.30 | 93.70 | 88.20 | 89.20 |
| **5→6** | 94.00 | 94.80 | 88.60 | 88.90 |
| **6→7** | 93.70 | 94.80 | 88.70 | 89.50 |
| **7→8** | 94.10 | 94.50 | 88.60 | 89.20 |
| **8→9** | 94.30 | 95.50 | 89.30 | 89.20 |
| **9→10** | 95.50 | 95.30 | 88.40 | 88.80 |

| | | | | |
|---|---|---|---|---|
| **10→11** | 95.60 | 95.30 | 88.50 | 88.10 |
| **11→12** | 95.40 | 95.20 | 87.90 | 87.60 |
| **12→13** | 95.50 | 96.00 | 86.90 | 86.90 |
| **13→14** | 96.20 | 96.20 | 85.20 | 85.60 |
| **14→15** | 96.70 | 96.10 | 86.60 | 86.10 |
| **15→16** | 96.70 | 96.90 | 84.80 | 84.60 |
| **16→17** | 96.90 | 96.90 | 83.20 | 83.20 |
| **17→18** | 96.70 | 96.50 | 84.30 | 83.80 |
| **18→19** | 97.30 | 96.90 | 82.10 | 82.30 |
| **19→20** | 95.80 | 96.30 | 81.90 | 81.30 |
| **20→21** | 95.50 | 96.20 | 81.40 | 81.80 |
| **21→22** | 95.00 | 95.50 | 80.90 | 80.40 |
| **22→23** | 91.30 | 93.50 | 80.10 | 79.80 |
| **23→24** | 78.60 | 87.70 | 80.20 | 80.20 |
| | **Beta-carbamate→intermediate-2** | | **intermediate-2→glutamate** | |
| | **Solvent** | **Complex** | **Solvent** | **Complex** |
| **λ** | **overlap (%)** | **overlap (%)** | **overlap (%)** | **overlap (%)** |
| **0→1** | 98.10 | 97.80 | 81.10 | 80.60 |
| **1→2** | 98.10 | 98.30 | 82.50 | 81.60 |
| **2→3** | 98.00 | 97.50 | 81.60 | 81.70 |
| **3→4** | 97.60 | 97.80 | 83.30 | 83.00 |

| | | | | |
|---|---|---|---|---|
| **4→5** | 97.60 | 97.60 | 82.50 | 82.40 |
| **5→6** | 97.00 | 97.50 | 84.80 | 84.40 |
| **6→7** | 97.40 | 97.00 | 85.00 | 84.50 |
| **7→8** | 97.00 | 97.30 | 85.60 | 86.80 |
| **8→9** | 97.20 | 96.80 | 87.10 | 86.80 |
| **9→10** | 95.30 | 97.00 | 87.20 | 86.80 |
| **10→11** | 96.20 | 96.50 | 87.80 | 87.30 |
| **11→12** | 95.60 | 96.40 | 89.50 | 89.30 |
| **12→13** | 95.60 | 95.60 | 89.80 | 89.70 |
| **13→14** | 94.40 | 96.10 | 89.00 | 89.40 |
| **14→15** | 94.40 | 95.40 | 91.10 | 91.00 |
| **15→16** | 94.30 | 94.80 | 90.70 | 91.00 |
| **16→17** | 94.90 | 95.00 | 92.20 | 92.50 |
| **17→18** | 95.00 | 94.50 | 91.20 | 91.70 |
| **18→19** | 93.30 | 94.60 | 93.20 | 93.00 |
| **19→20** | 93.30 | 94.20 | 91.00 | 92.00 |
| **20→21** | 93.10 | 92.80 | 92.60 | 92.70 |
| **21→22** | 93.40 | 93.40 | 91.00 | 91.40 |
| **22→23** | 93.30 | 88.30 | 90.10 | 89.80 |
| **23→24** | 93.10 | 94.80 | 87.70 | 85.30 |

**Table S7.** Phase space overlap between neighboring λ windows of perturbations glutamate→intermediate-1, intermediate-1→beta-carbamate, beta-carbamate→intermediate-2, and intermediate-2→glutamate. Results obtained from replica 3 of the AMBER20/ff14SB/TI protocol.

| | **Glutamate→intermediate-1** | | **intermediate-1→beta-carbamate** | |
|---|---|---|---|---|
| | **Solvent** | **Complex** | **Solvent** | **Complex** |
| **λ** | **overlap (%)** | **overlap (%)** | **overlap (%)** | **overlap (%)** |
| **0→1** | 94.00 | 93.20 | 21.20 | 20.00 |
| **1→2** | 93.90 | 93.90 | 73.30 | 75.70 |
| **2→3** | 93.30 | 94.10 | 84.50 | 84.50 |
| **3→4** | 93.90 | 93.90 | 85.60 | 87.40 |
| **4→5** | 93.60 | 94.50 | 87.90 | 88.70 |
| **5→6** | 93.80 | 94.70 | 88.60 | 89.30 |
| **6→7** | 94.10 | 95.00 | 88.80 | 89.60 |
| **7→8** | 93.90 | 94.60 | 88.80 | 89.20 |
| **8→9** | 94.80 | 94.90 | 89.40 | 89.10 |
| **9→10** | 95.40 | 95.00 | 88.30 | 88.60 |
| **10→11** | 95.00 | 95.50 | 88.20 | 88.60 |
| **11→12** | 95.30 | 95.30 | 87.70 | 87.70 |
| **12→13** | 96.00 | 95.70 | 86.90 | 86.90 |
| **13→14** | 96.20 | 95.90 | 85.50 | 85.40 |

| | | | | |
|---|---|---|---|---|
| **14→15** | 96.10 | 96.40 | 86.70 | 85.90 |
| **15→16** | 96.80 | 97.00 | 84.40 | 84.90 |
| **16→17** | 96.50 | 96.80 | 82.80 | 83.30 |
| **17→18** | 96.80 | 96.70 | 84.40 | 83.90 |
| **18→19** | 96.70 | 96.50 | 82.10 | 81.70 |
| **19→20** | 96.60 | 96.70 | 81.60 | 81.50 |
| **20→21** | 95.70 | 96.20 | 81.50 | 81.50 |
| **21→22** | 94.40 | 94.70 | 80.50 | 80.50 |
| **22→23** | 91.80 | 94.20 | 80.00 | 80.50 |
| **23→24** | 77.30 | 87.00 | 80.40 | 79.80ß |
| | **Beta-carbamate→intermediate-2** | | **intermediate-2→glutamate** | |
| | **Solvent** | **Complex** | **Solvent** | **Complex** |
| **λ** | **overlap (%)** | **overlap (%)** | **overlap (%)** | **overlap (%)** |
| **0→1** | 98.10 | 98.10 | 81.10 | 81.20 |
| **1→2** | 98.30 | 98.10 | 81.80 | 81.50 |
| **2→3** | 98.20 | 97.60 | 81.70 | 81.90 |
| **3→4** | 97.70 | 97.60 | 83.90 | 82.10 |
| **4→5** | 97.50 | 97.20 | 82.10 | 82.20 |
| **5→6** | 97.50 | 97.40 | 85.10 | 84.80 |
| **6→7** | 96.90 | 97.40 | 84.60 | 84.10 |

| | | | | |
|---|---|---|---|---|
| **7→8** | 97.10 | 96.90 | 85.30 | 84.70 |
| **8→9** | 96.60 | 97.30 | 87.50 | 86.50 |
| **9→10** | 96.20 | 96.80 | 87.00 | 86.40 |
| **10→11** | 96.70 | 96.20 | 87.20 | 87.50 |
| **11→12** | 95.50 | 96.50 | 89.70 | 89.10 |
| **12→13** | 94.40 | 96.50 | 90.30 | 89.60 |
| **13→14** | 94.70 | 95.90 | 89.00 | 89.30 |
| **14→15** | 94.40 | 95.70 | 90.90 | 91.10 |
| **15→16** | 94.90 | 95.80 | 90.70 | 90.90 |
| **16→17** | 94.90 | 95.50 | 92.40 | 92.60 |
| **17→18** | 94.40 | 95.70 | 91.10 | 92.00 |
| **18→19** | 93.30 | 95.10 | 92.80 | 93.60 |
| **19→20** | 93.20 | 93.00 | 91.30 | 92.10 |
| **20→21** | 93.30 | 94.50 | 92.60 | 93.60 |
| **21→22** | 93.70 | 90.60 | 91.30 | 92.30 |
| **22→23** | 93.40 | 93.70 | 90.40 | 92.20 |
| **23→24** | 93.30 | 92.40 | 86.80 | 89.20 |

**FIGURES**

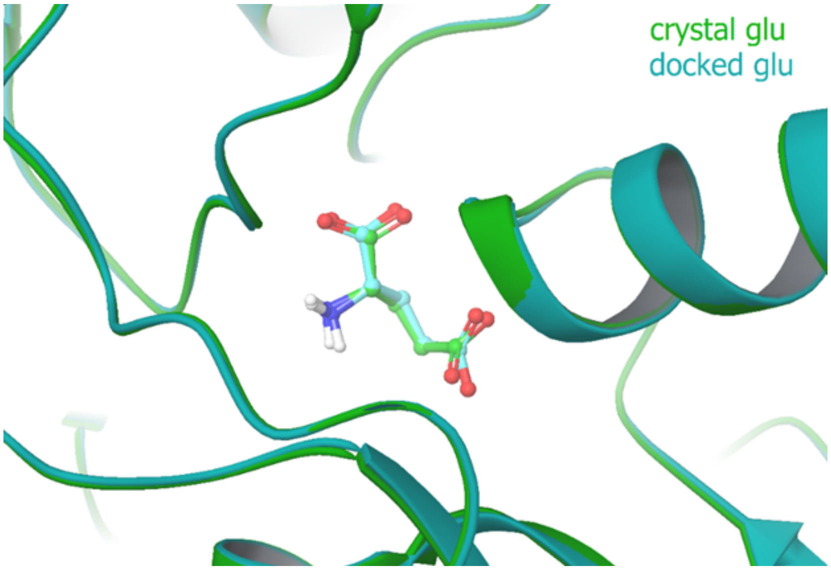


**Figure S1.** Alignment between the crystal of glutamate (colored in green) in PDB ID 1FTJ and docked glutamate (colored in blue).

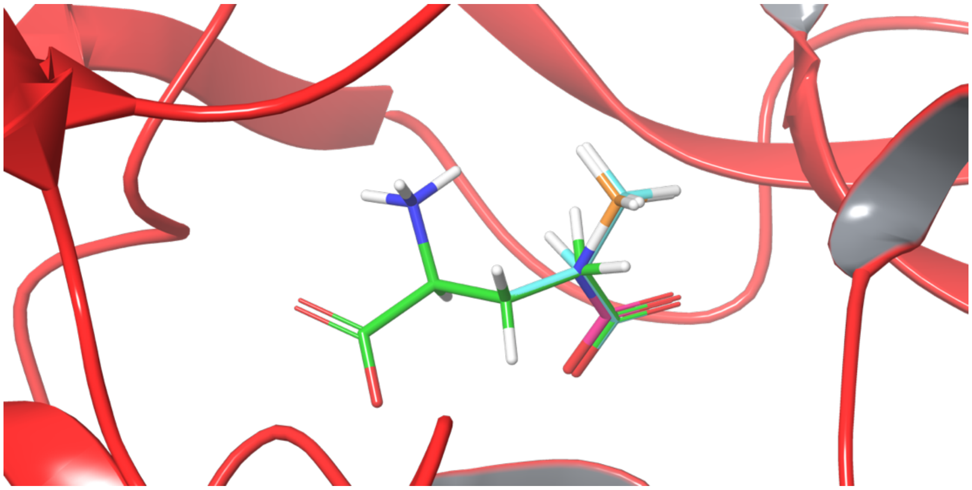

**Figure S2.** The 3D chemical structures of glutamate (colored in green), intermediate-1 (colored in magenta), intermediate-2 (colored in light blue), and beta-carbamate (colored in orange).

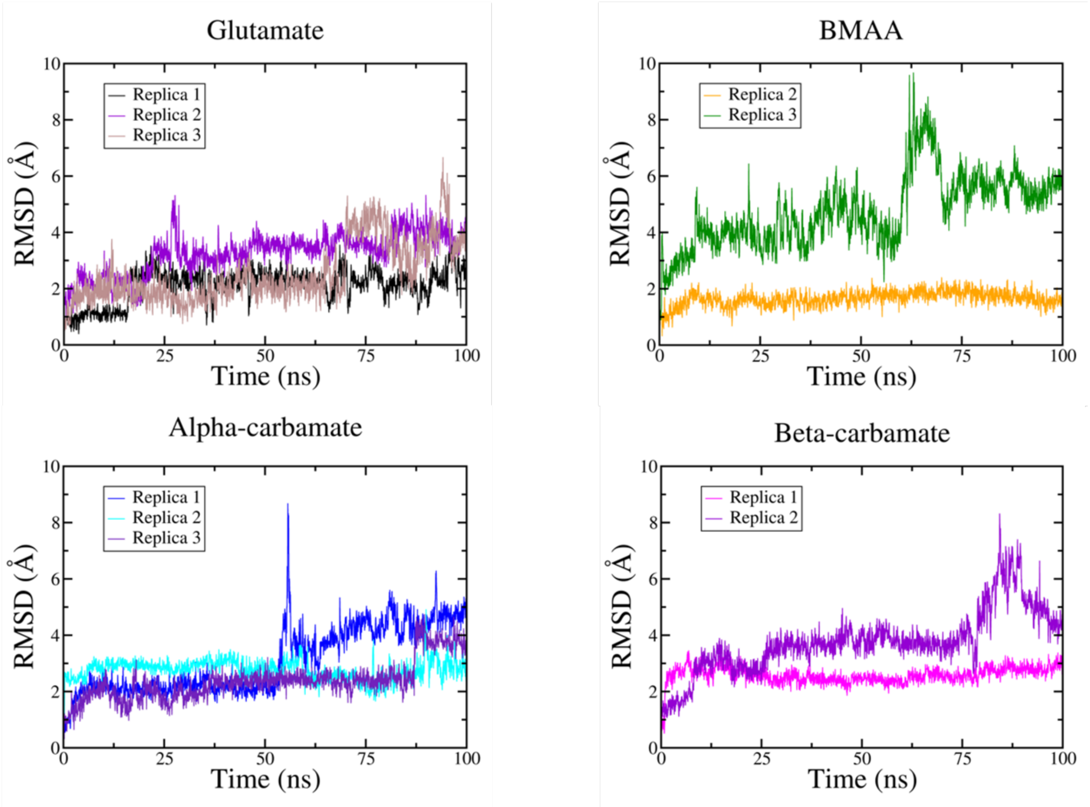


**Figure S3.** RMSD of the three replicas of glutamate, BMAA, alpha-carbamate, and beta-carbamate as a function of time using NAMD2.14/OPLS-AA.

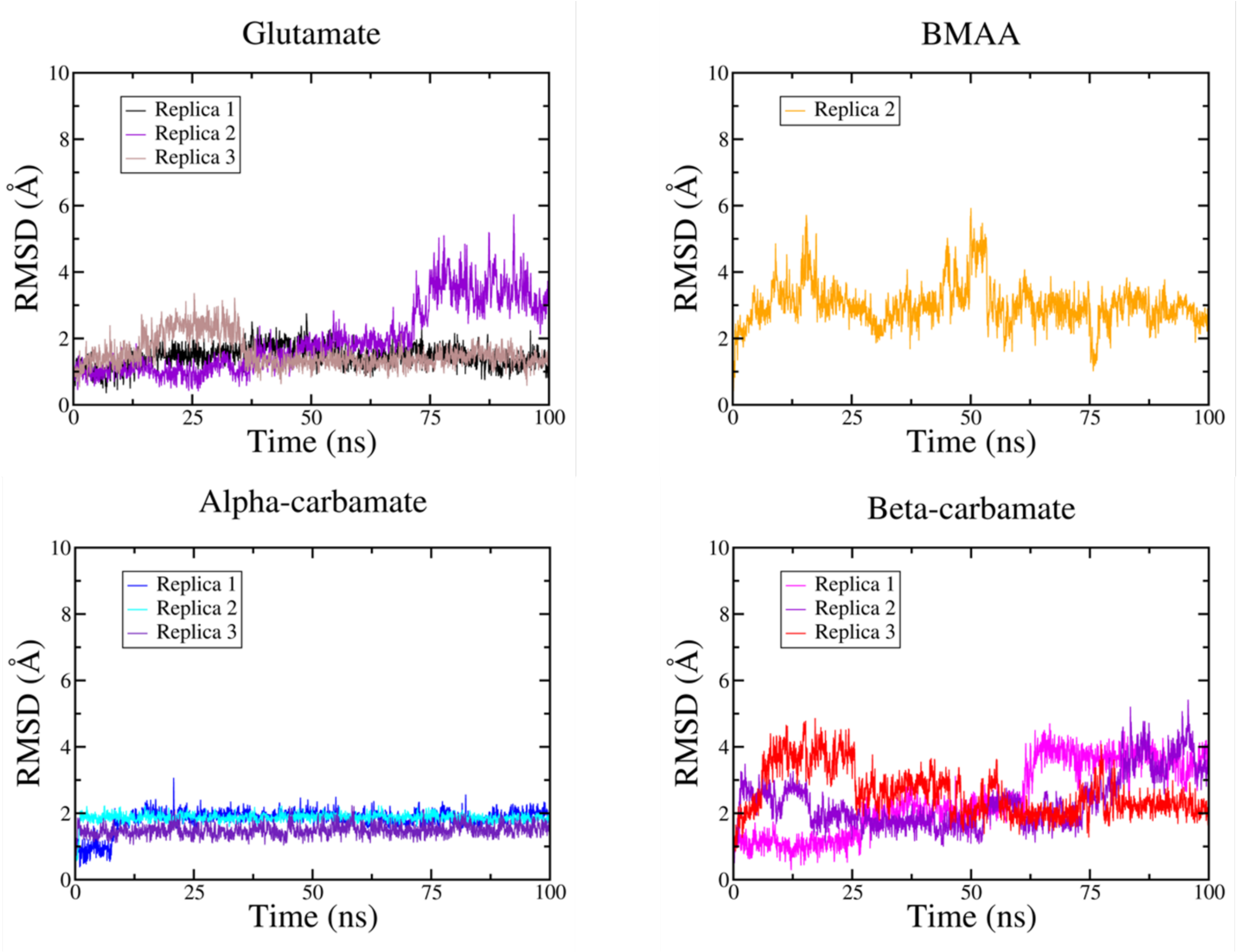


**Figure S4.** RMSD of the three replicas of glutamate, BMAA, alpha-carbamate, and beta-carbamate as a function of time using AMBER20/ff19SB.

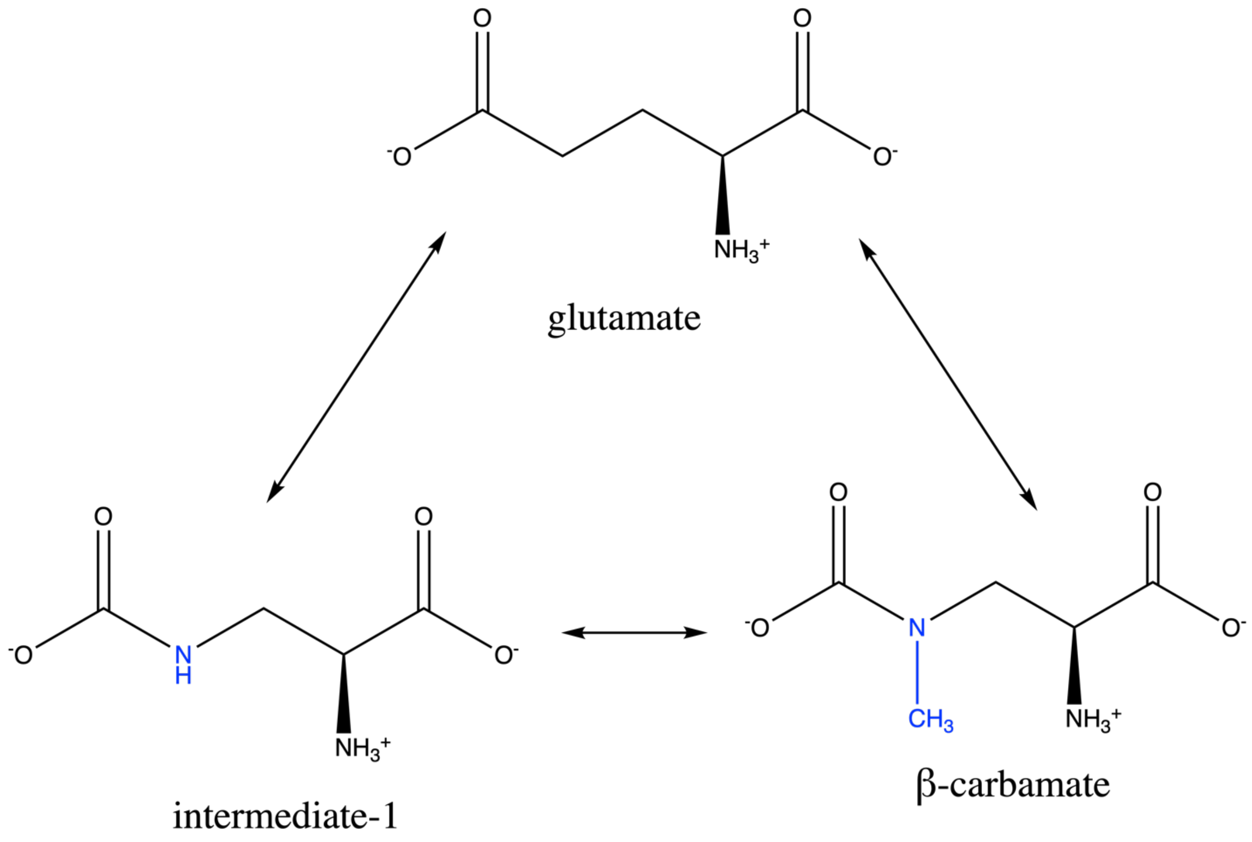


**Figure S5.** Perturbation network used for the initial RBFE calculations. Glutamate and beta-carbamate are connected using one intermediate molecule. The perturbed atoms with respect to glutamate are coloured blue.

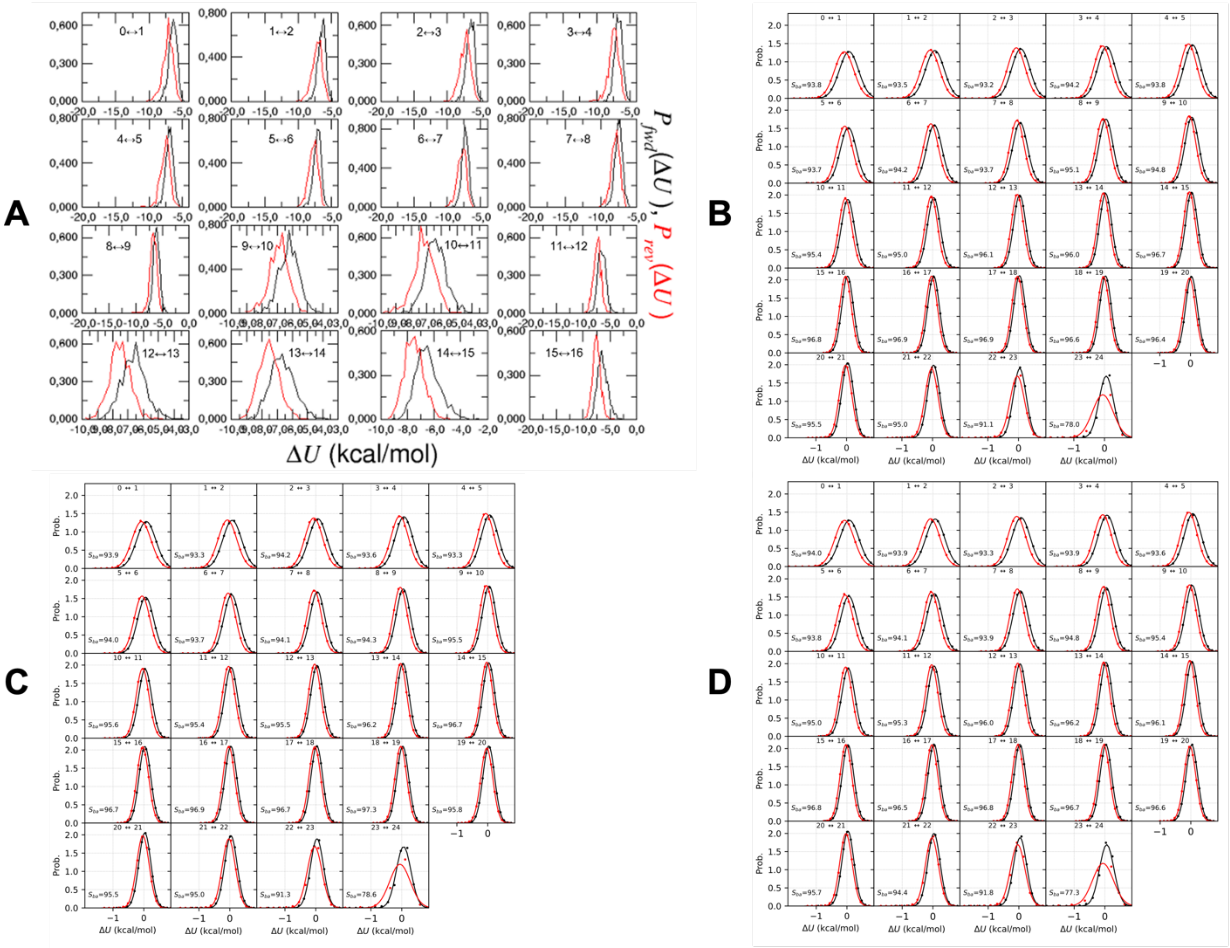

**Figure S6.** Histograms of the probability distribution of the forward, $P_{fwd}(\Delta U)$ (black solid line) and backward, $P_{rev}(\Delta U)$ (red solid line) calculations for the perturbation of glutamate to intermediate-1 in the solvent phase obtained from A) NAMD2.14/OPLS-AA/FEP protocol, B) replica1 of the AMBER20/ff14SB/TI protocol, C) replica2 of the AMBER20/ff14SB/TI protocol, and D) replica3 of the AMBER20/ff14SB/TI protocol.

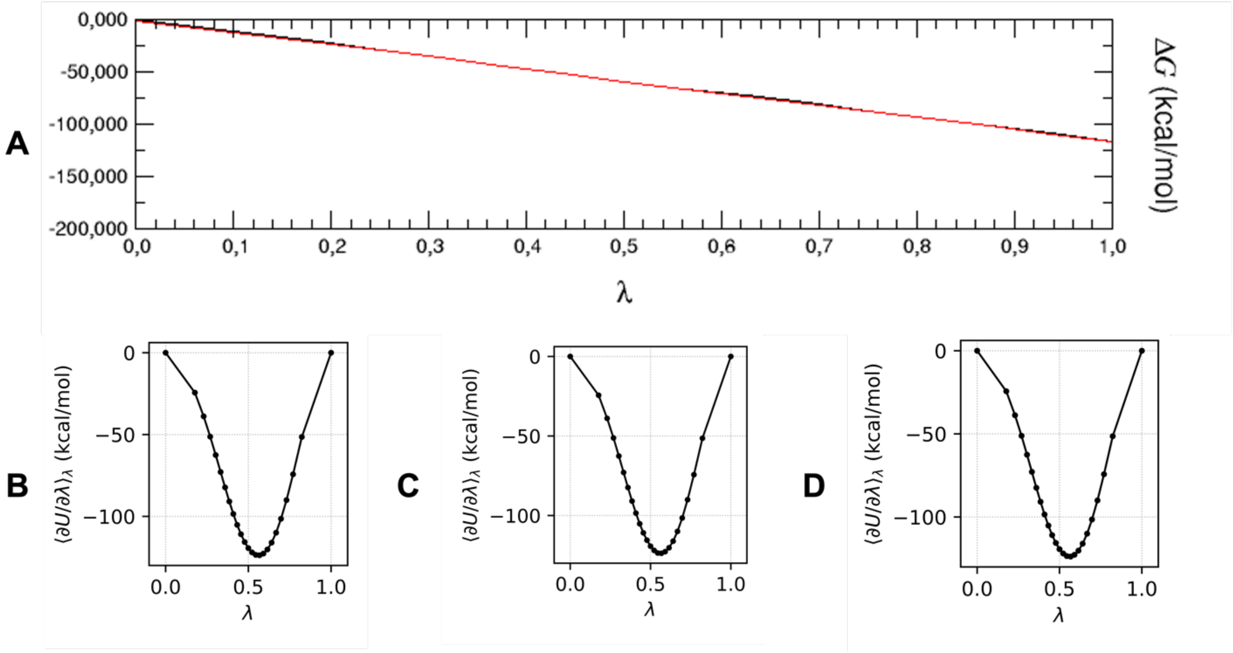


**Figure S7.** A) Free energy (ΔG) versus the coupling parameter λ for the forward (black solid line) and backward (red solid line) perturbation of glutamate to intermediate-1 in the solvent phase obtained from NAMD2.14/OPLS-AA/FEP protocol. Variation of the gradient of the potential energy (∂U/∂λ) as a function of λ versus the coupling parameter λ for the perturbation of glutamate to intermediate-1in the solvent phase obtained from B) replica1 of the AMBER20/ff14SB/TI protocol, C) replica2 of the AMBER20/ff14SB/TI protocol, and D) replica3 of the AMBER20/ff14SB/TI protocol

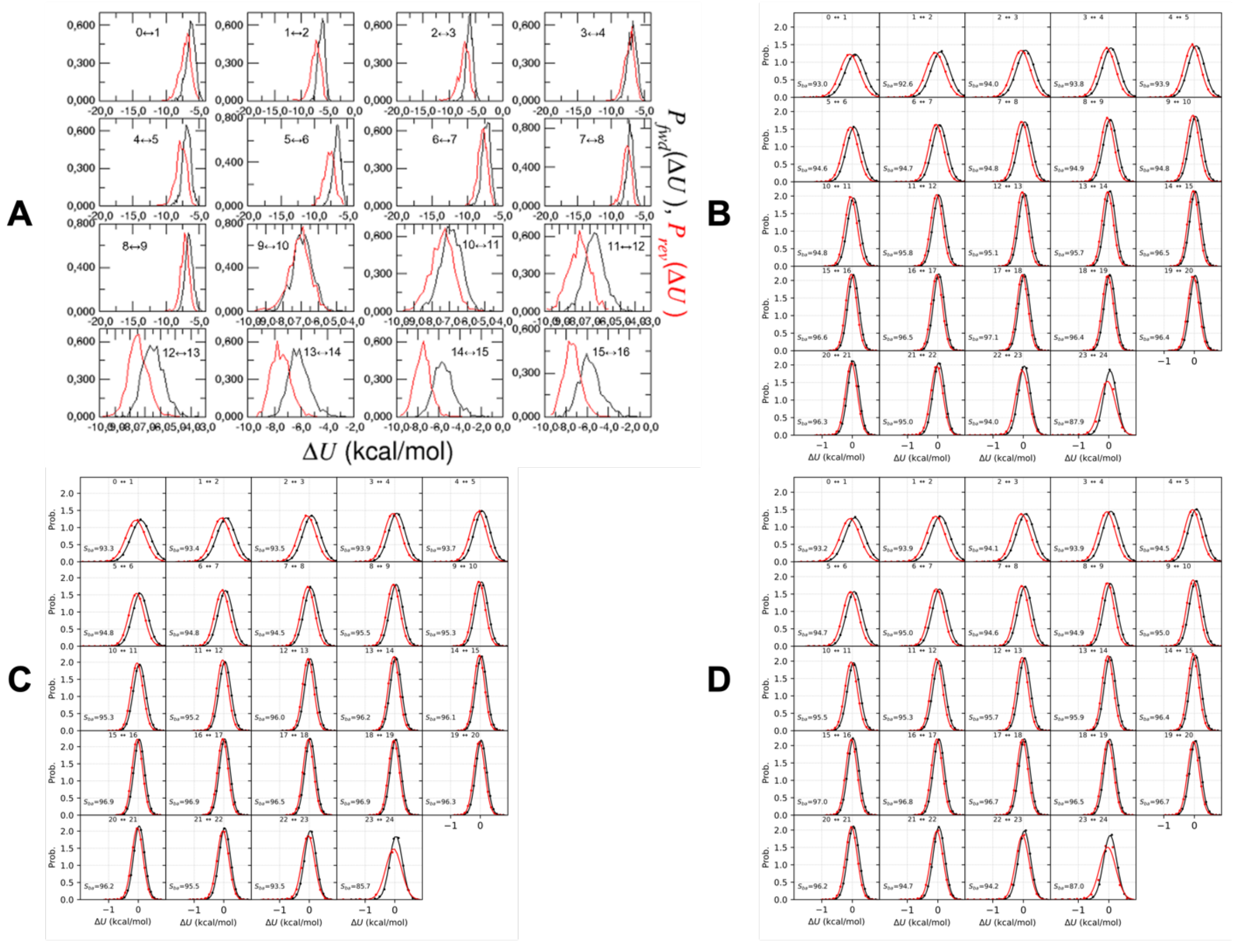


**Figure S8.** Histograms of the probability distribution of the forward, $P_{fwd}(\Delta U)$ (black solid line) and backward, $P_{rev}(\Delta U)$ (red solid line) calculations for the perturbation of glutamate to intermediate-1 in the complex phase obtained from A) NAMD2.14/OPLS-AA/FEP protocol, B) replica1 of the AMBER20/ff14SB/TI protocol, C) replica2 of the AMBER20/ff14SB/TI protocol, and D) replica3 of the AMBER20/ff14SB/TI protocol

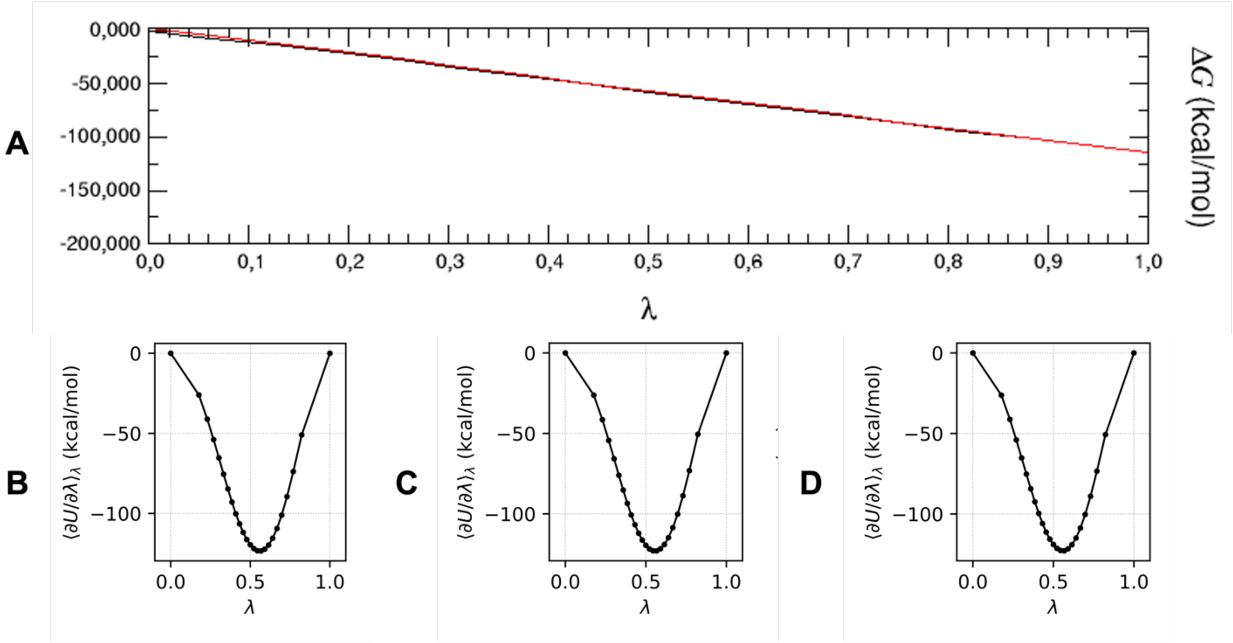


**Figure S9.** A) Free energy (ΔG) versus the coupling parameter λ for the forward (black solid line) and backward (red solid line) perturbation of glutamate to intermediate-1 in the complex phase obtained from NAMD2.14/OPLS-AA/FEP protocol. Variation of the gradient of the potential energy (∂U/∂λ) as a function of λ versus the coupling parameter λ for the perturbation of glutamate to intermediate-1in the complex phase obtained from B) replica1 of the AMBER20/ff14SB/TI protocol, C) replica2 of the AMBER20/ff14SB/TI protocol, and D) replica3 of the AMBER20/ff14SB/TI protocol

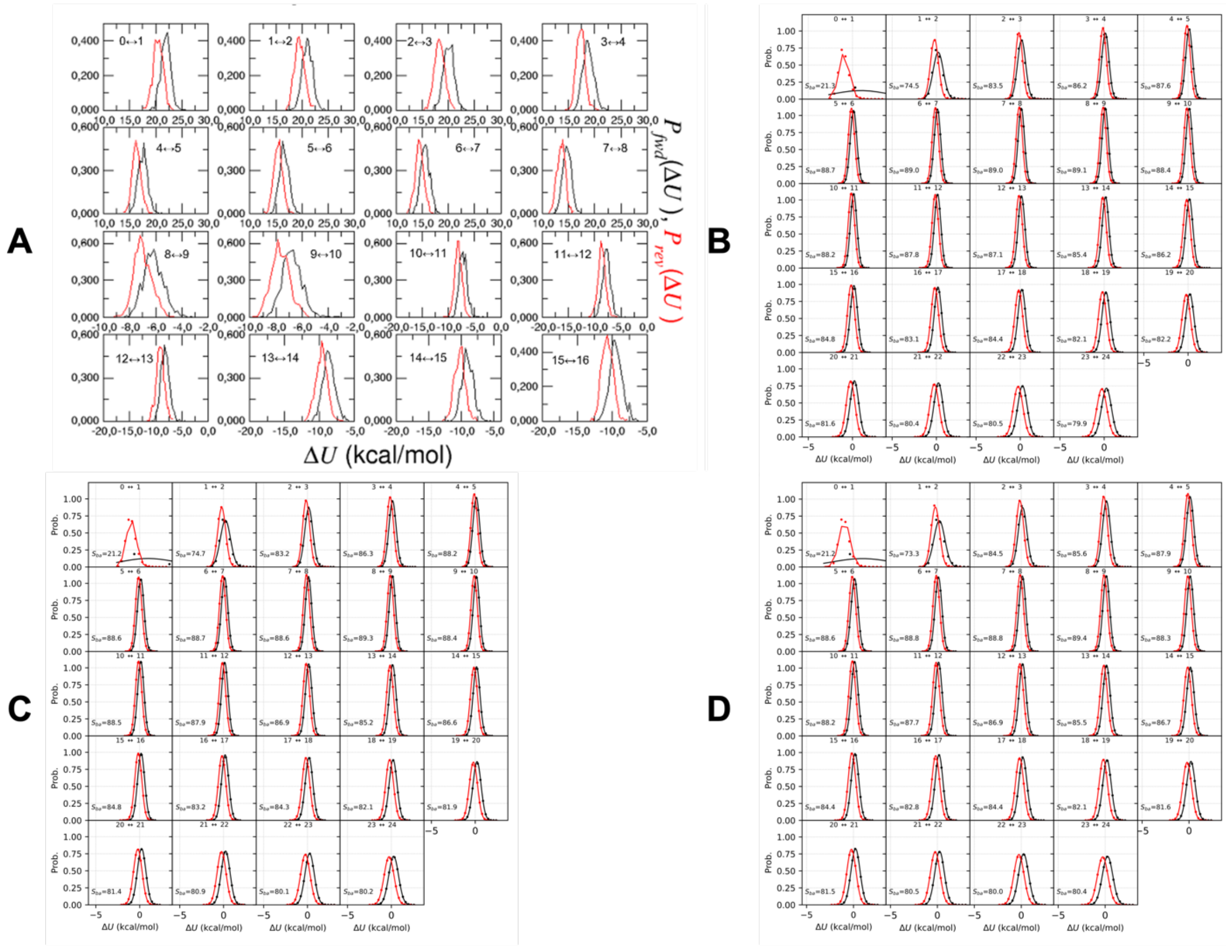

**Figure S10.** Histograms of the probability distribution of the forward, $P_{fwd}(\Delta U)$ (black solid line) and backward, $P_{rev}(\Delta U)$ (red solid line) calculations for the perturbation of intermediate-1 to beta-carbamate in the solvent phase obtained from A) NAMD2.14/OPLS-AA/FEP protocol, B) replica1 of the AMBER20/ff14SB/TI protocol, C) replica2 of the AMBER20/ff14SB/TI protocol, and D) replica3 of the AMBER20/ff14SB/TI protocol

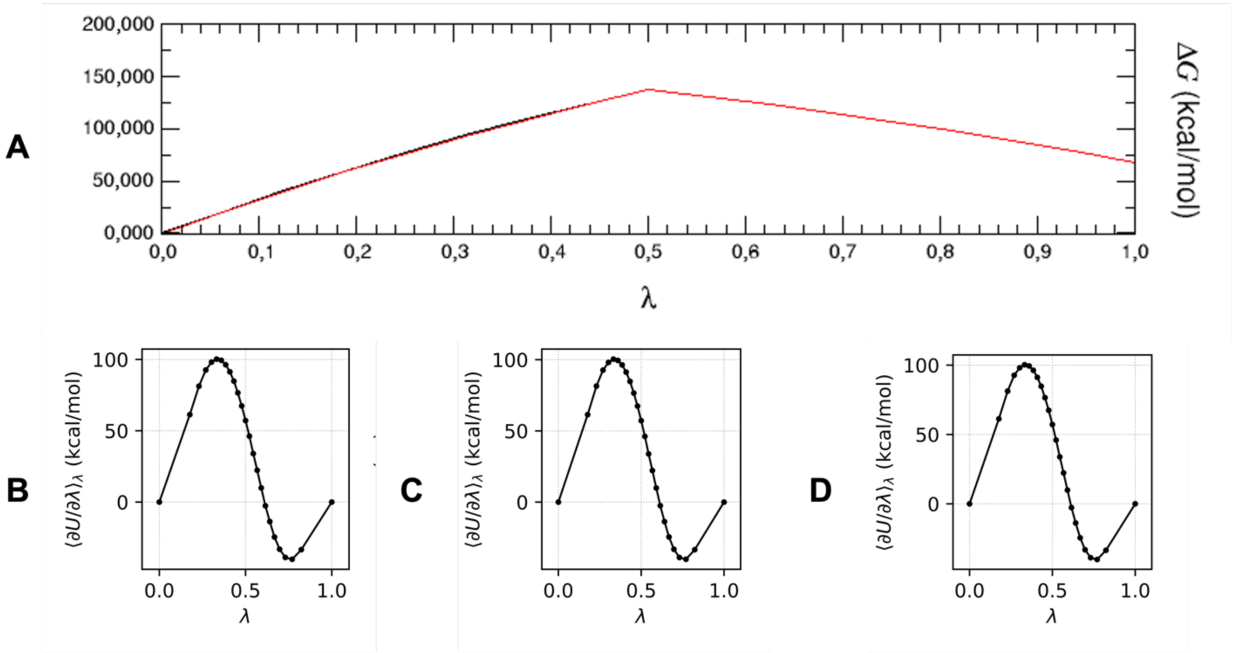


**Figure S11.** A) Free energy (ΔG) versus the coupling parameter λ for the forward (black solid line) and backward (red solid line) perturbation of intermediate-1 to beta-carbamate in the solvent phase obtained from NAMD2.14/OPLS-AA/FEP protocol. Variation of the gradient of the potential energy (∂U/∂λ) as a function of λ versus the coupling parameter λ for the perturbation of intermediate-1 to beta-carbamate in the solvent phase obtained from B) replica1 of the AMBER20/ff14SB/TI protocol, C) replica2 of the AMBER20/ff14SB/TI protocol, and D) replica3 of the AMBER20/ff14SB/TI protocol

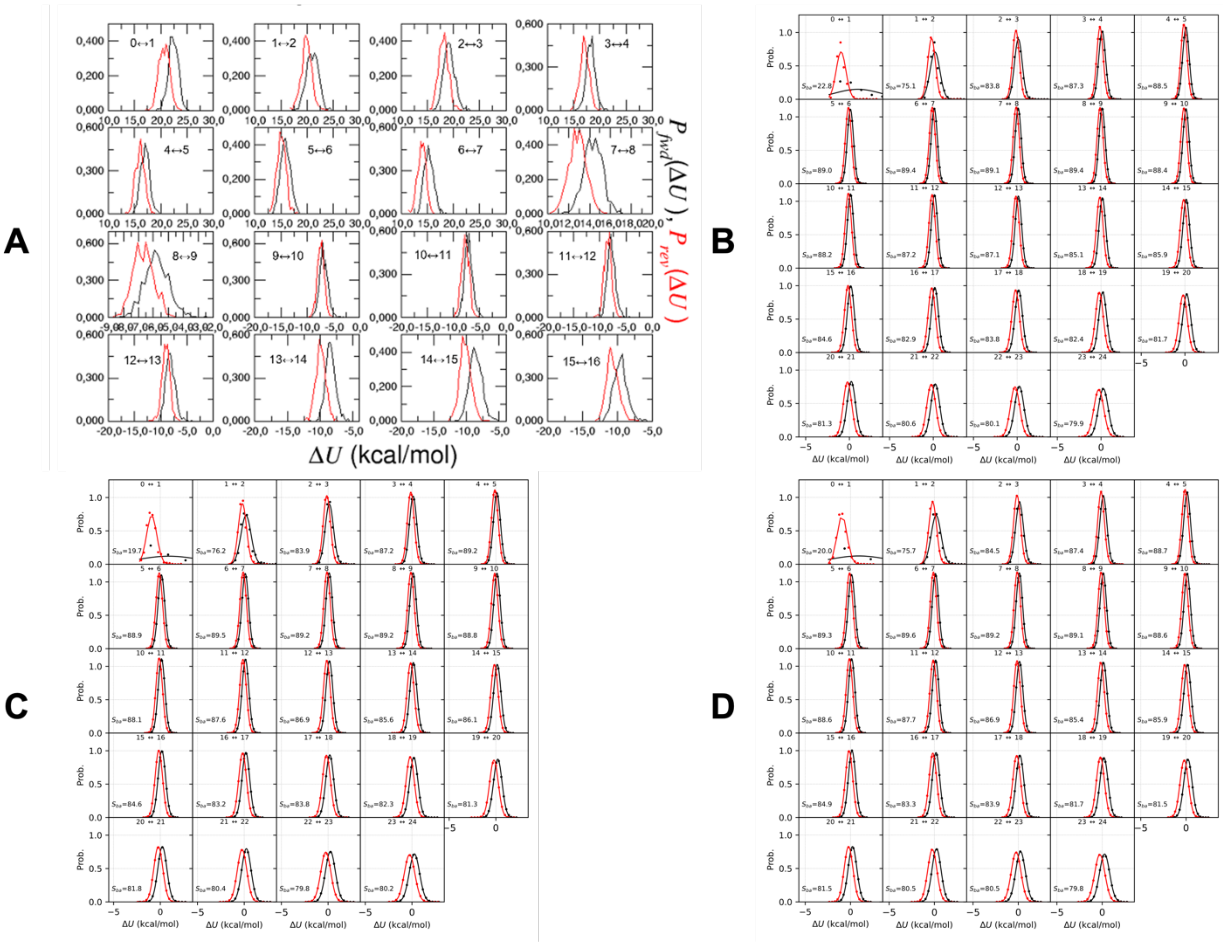


**Figure S12.** Histograms of the probability distribution of the forward, $P_{fwd}(\Delta U)$ (black solid line) and backward, $P_{rev}(\Delta U)$ (red solid line) calculations for the perturbation of intermediate-1 to beta-carbamate in the complex phase obtained from A) NAMD2.14/OPLS-AA/FEP protocol, B) replica1 of the AMBER20/ff14SB/TI protocol, C) replica2 of the AMBER20/ff14SB/TI protocol, and D) replica3 of the AMBER20/ff14SB/TI protocol

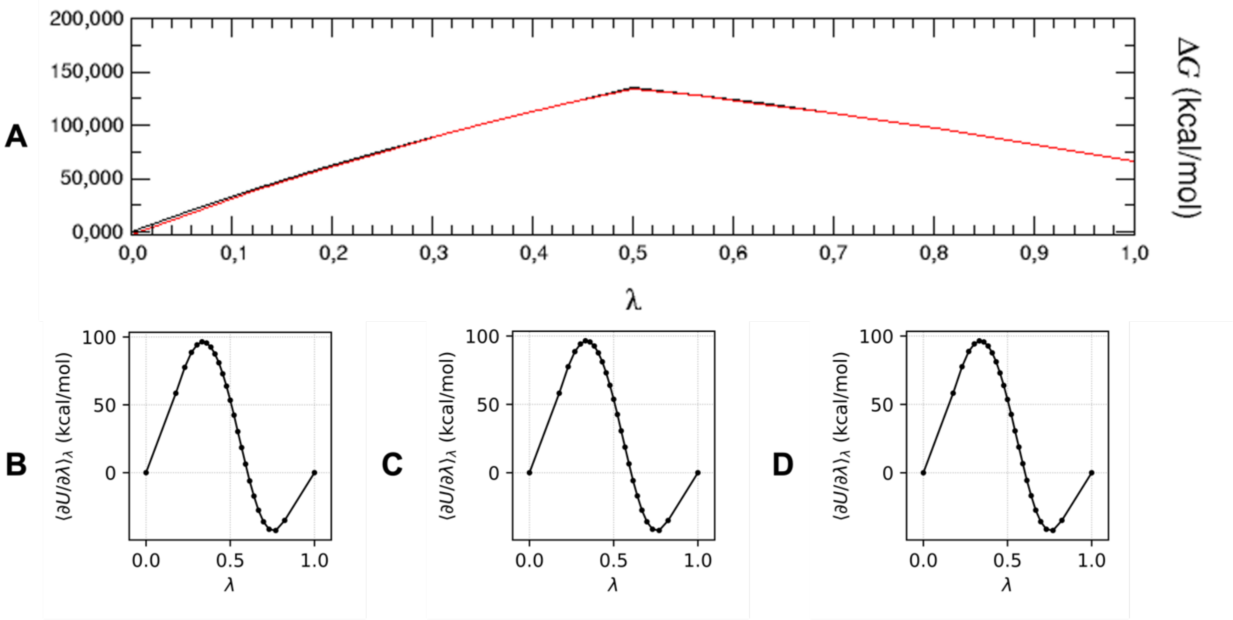


**Figure S13.** A) Free energy (ΔG) versus the coupling parameter λ for the forward (black solid line) and backward (red solid line) perturbation of intermediate-1 to beta-carbamate in the complex phase obtained from NAMD2.14/OPLS-AA/FEP protocol. Variation of the gradient of the potential energy (∂U/∂λ) as a function of λ versus the coupling parameter λ for the perturbation of intermediate-1 to beta-carbamate in the complex phase obtained from B) replica1 of the AMBER20/ff14SB/TI protocol, C) replica2 of the AMBER20/ff14SB/TI protocol, and D) replica3 of the AMBER20/ff14SB/TI protocol

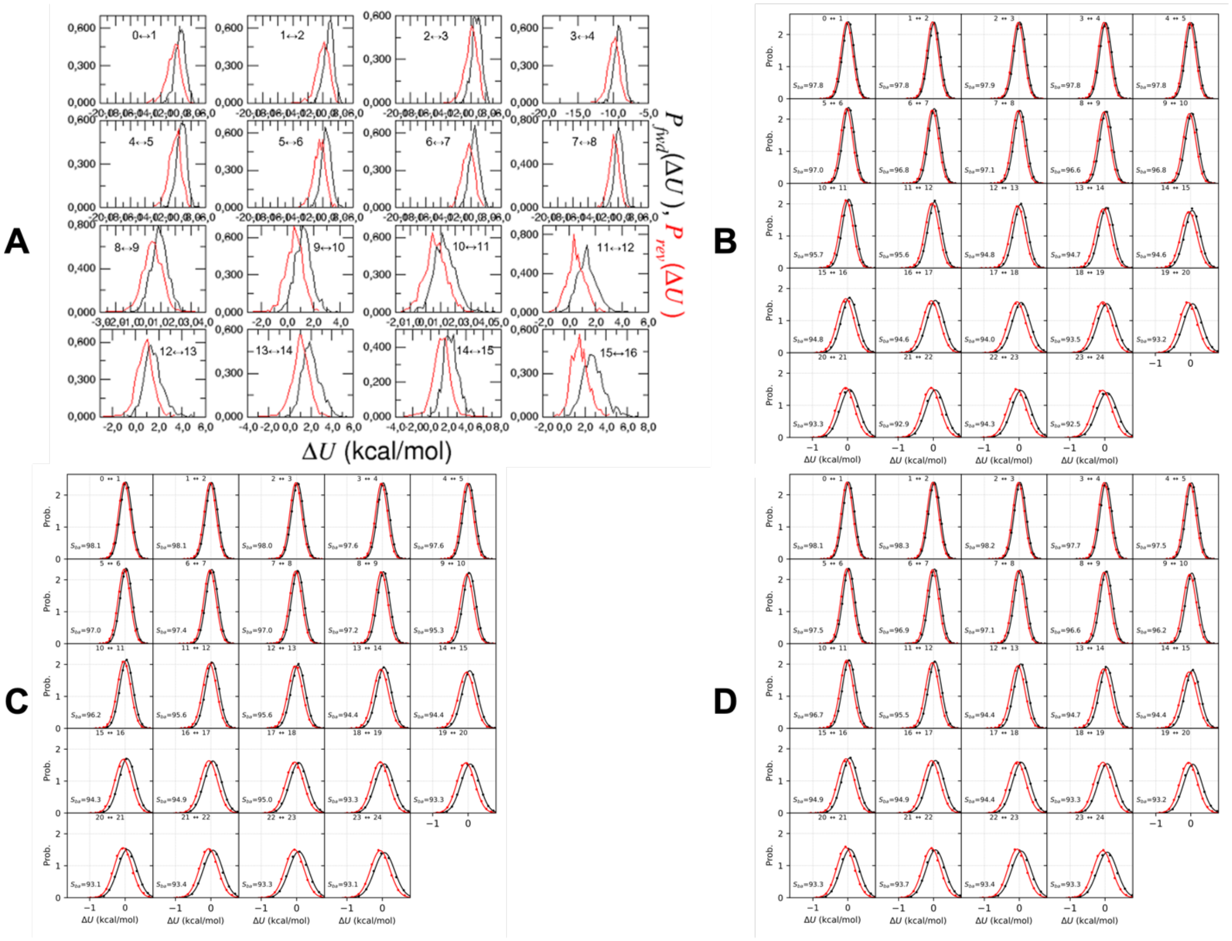


**Figure S14.** Histograms of the probability distribution of the forward, $P_{fwd}(\Delta U)$ (black solid line) and backward, $P_{rev}(\Delta U)$ (red solid line) calculations for the perturbation of beta-carbamate to intermediate-2 in the solvent phase obtained from A) NAMD2.14/OPLS-AA/FEP protocol, B) replica1 of the AMBER20/ff14SB/TI protocol, C) replica2 of the AMBER20/ff14SB/TI protocol, and D) replica3 of the AMBER20/ff14SB/TI protocol

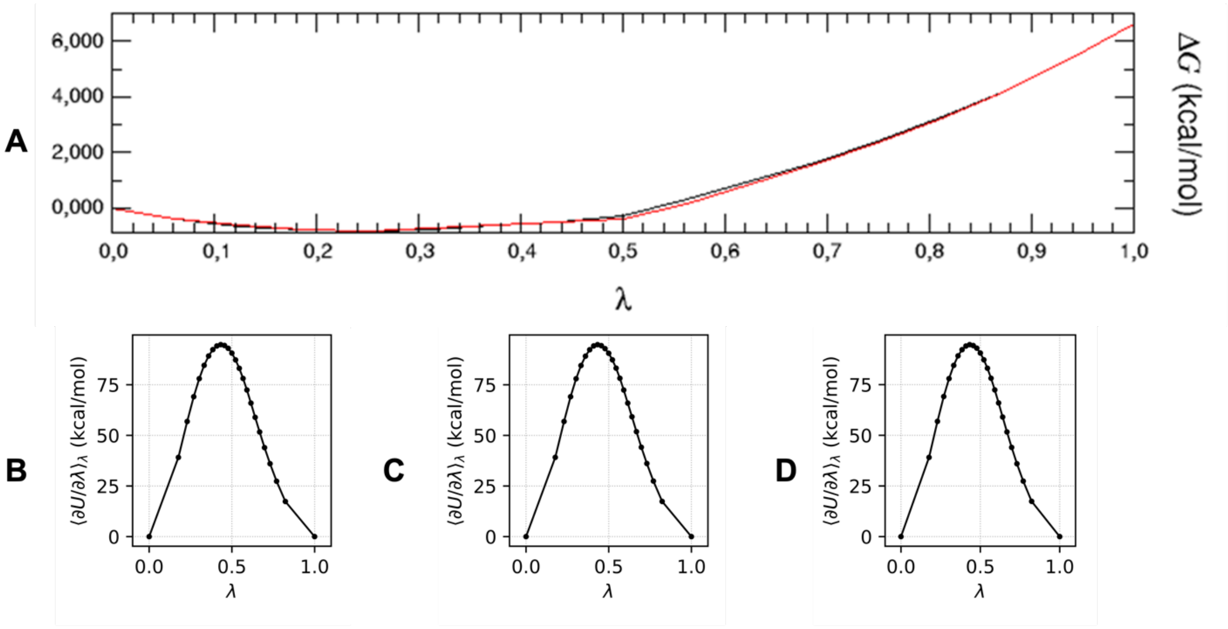


**Figure S15.** A) Free energy (ΔG) versus the coupling parameter λ for the forward (black solid line) and backward (red solid line) perturbation of beta-carbamate to intermediate-2 in the solvent phase obtained from NAMD2.14/OPLS-AA/FEP protocol. Variation of the gradient of the potential energy (∂U/∂λ) as a function of λ versus the coupling parameter λ for the perturbation of intermediate-2 to glutamate in the solvent phase obtained from B) replica1 of the AMBER20/ff14SB/TI protocol, C) replica2 of the AMBER20/ff14SB/TI protocol, and D) replica3 of the AMBER20/ff14SB/TI protocol

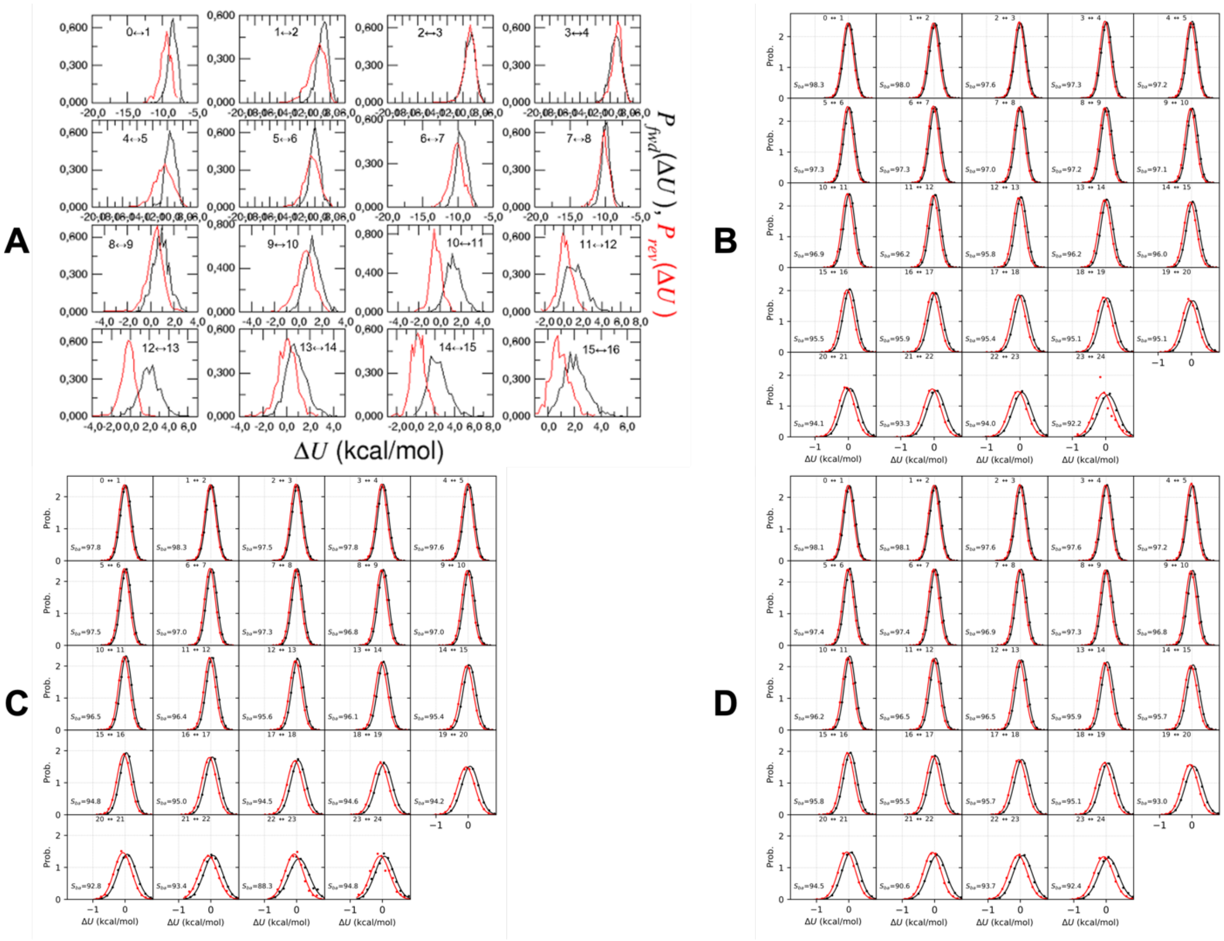


**Figure S16.** Histograms of the probability distribution of the forward, $P_{fwd}(\Delta U)$ (black solid line) and backward, $P_{rev}(\Delta U)$ (red solid line) calculations for the perturbation of beta-carbamate to intermediate-2 in the complex phase obtained from A) NAMD2.14/OPLS-AA/FEP protocol, B) replica1 of the AMBER20/ff14SB/TI protocol, C) replica2 of the AMBER20/ff14SB/TI protocol, and D) replica3 of the AMBER20/ff14SB/TI protocol

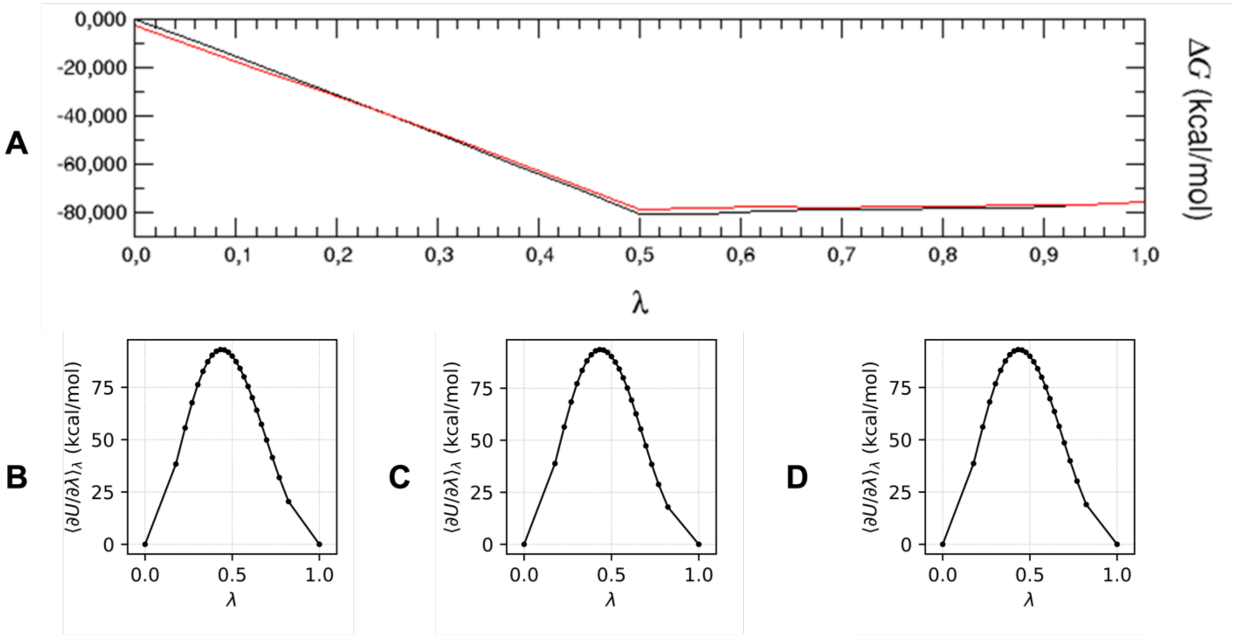


**Figure S17.** A) Free energy (ΔG) versus the coupling parameter λ for the forward (black solid line) and backward (red solid line) perturbation of beta-carbamate to intermediate-2 in the complex phase obtained from NAMD2.14/OPLS-AA/FEP protocol. Variation of the gradient of the potential energy (∂U/∂λ) as a function of λ versus the coupling parameter λ for the perturbation of intermediate-2 to glutamate in the complex phase obtained from B) replica1 of the AMBER20/ff14SB/TI protocol, C) replica2 of the AMBER20/ff14SB/TI protocol, and D) replica3 of the AMBER20/ff14SB/TI protocol

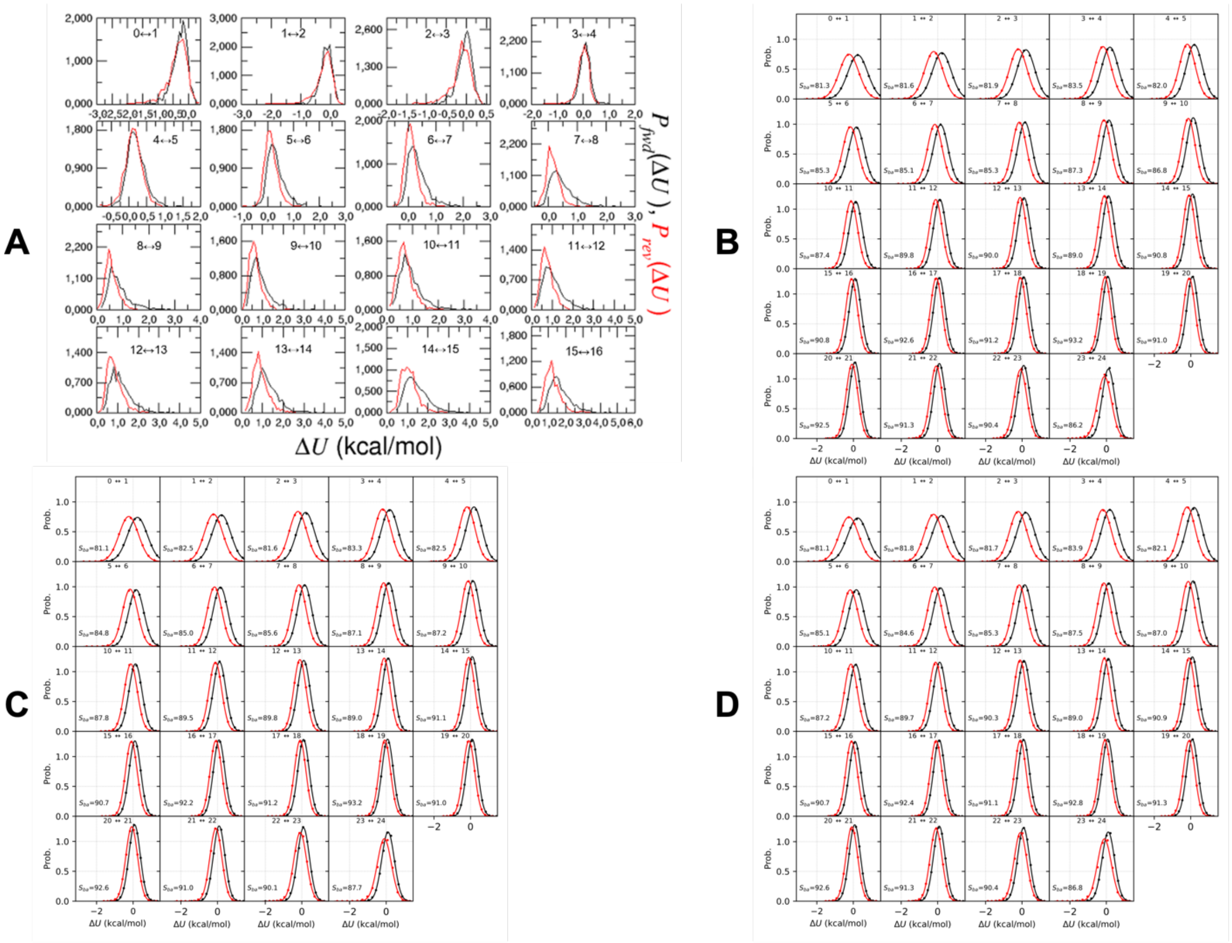


**Figure S18.** Histograms of the probability distribution of the forward, $P_{fwd}(\Delta U)$ (black solid line) and backward, $P_{rev}(\Delta U)$ (red solid line) calculations for the perturbation of intermediate-2 to glutamate in the solvent phase obtained from A) NAMD2.14/OPLS-AA/FEP protocol, B) replica1 of the AMBER20/ff14SB/TI protocol, C) replica2 of the AMBER20/ff14SB/TI protocol, and D) replica3 of the AMBER20/ff14SB/TI protocol

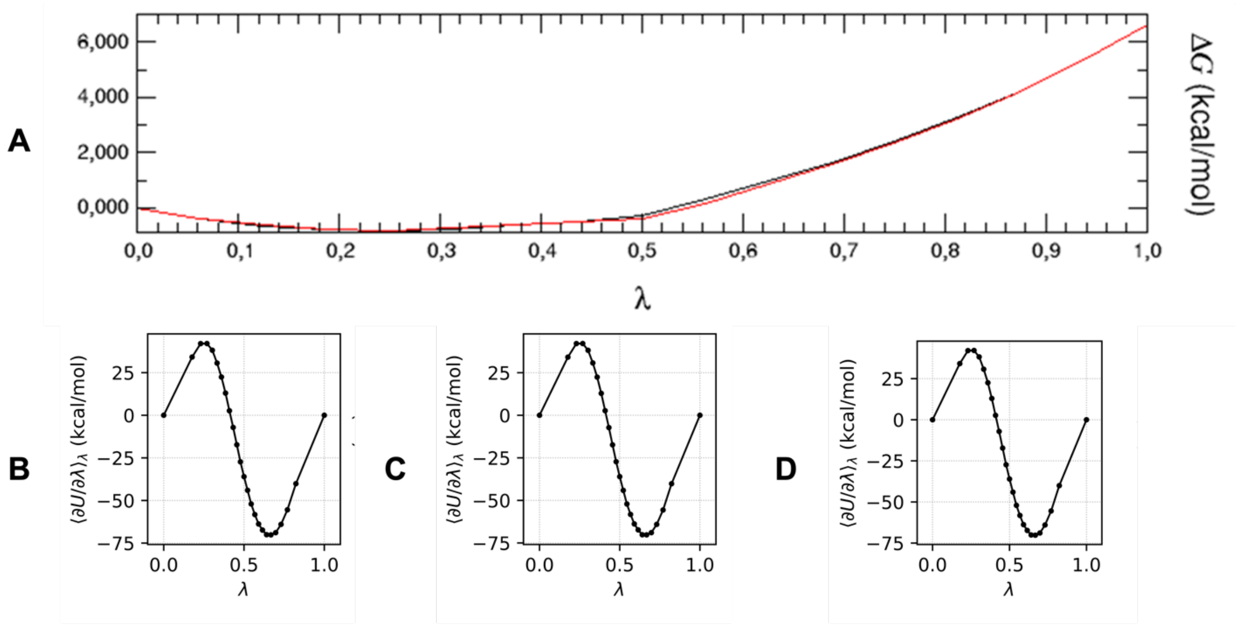


**Figure S19.** A) Free energy (ΔG) versus the coupling parameter λ for the forward (black solid line) and backward (red solid line) perturbation of intermediate-2 to glutamate in the solvent phase obtained from NAMD2.14/OPLS-AA/FEP protocol. Variation of the gradient of the potential energy (∂U/∂λ) as a function of λ versus the coupling parameter λ for the perturbation of intermediate-2 to glutamate in the solvent phase obtained from B) replica1 of the AMBER20/ff14SB/TI protocol, C) replica2 of the AMBER20/ff14SB/TI protocol, and D) replica3 of the AMBER20/ff14SB/TI protocol

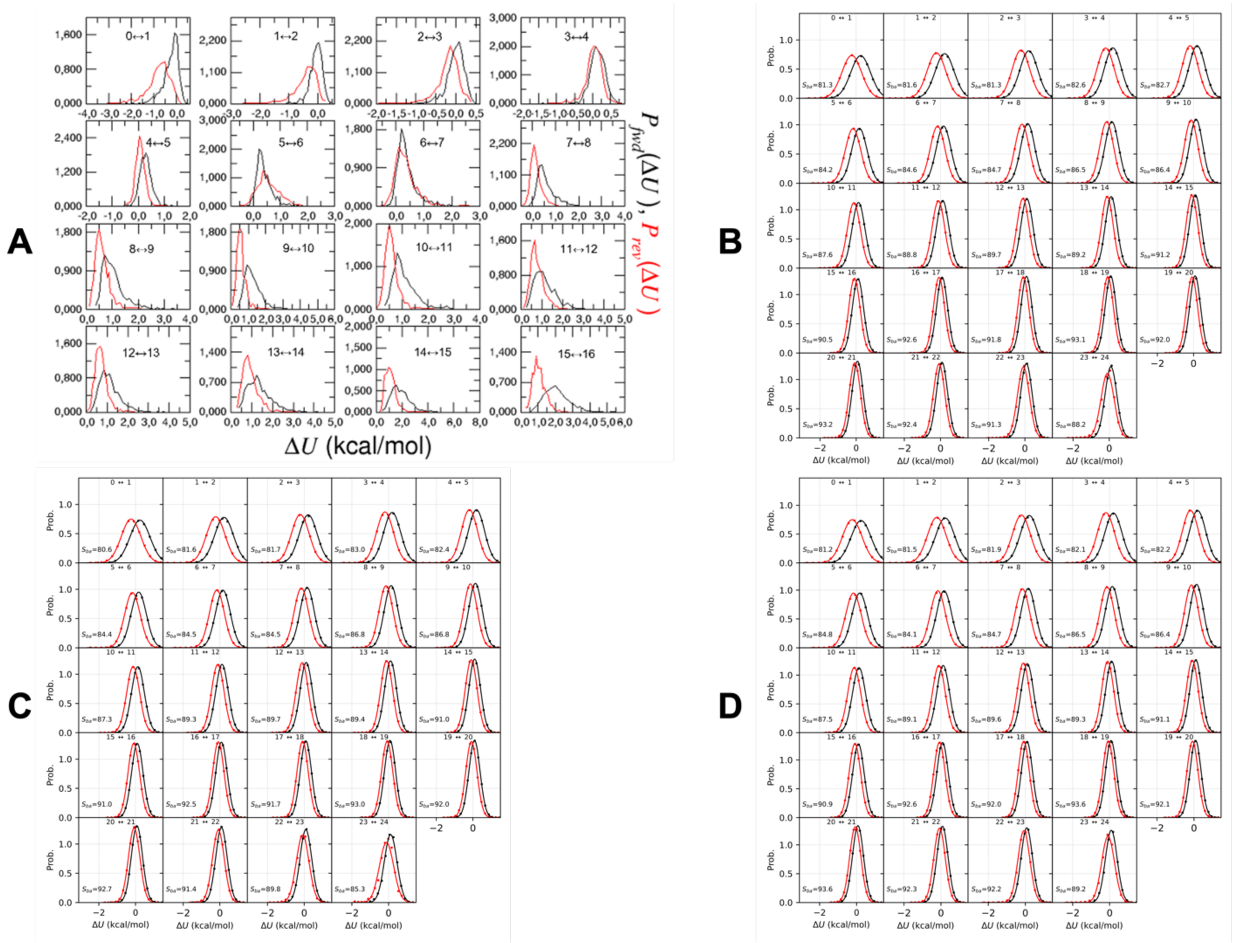


**Figure S20.** Histograms of the probability distribution of the forward, $P_{fwd}(\Delta U)$ (black solid line) and backward, $P_{rev}(\Delta U)$ (red solid line) calculations for the perturbation of intermediate-2 to glutamate in complex phase obtained from A) NAMD2.14/OPLS-AA/FEP protocol, B) replica1 of the AMBER20/ff14SB/TI protocol, C) replica2 of the AMBER20/ff14SB/TI protocol, and D) replica3 of the AMBER20/ff14SB/TI protocol

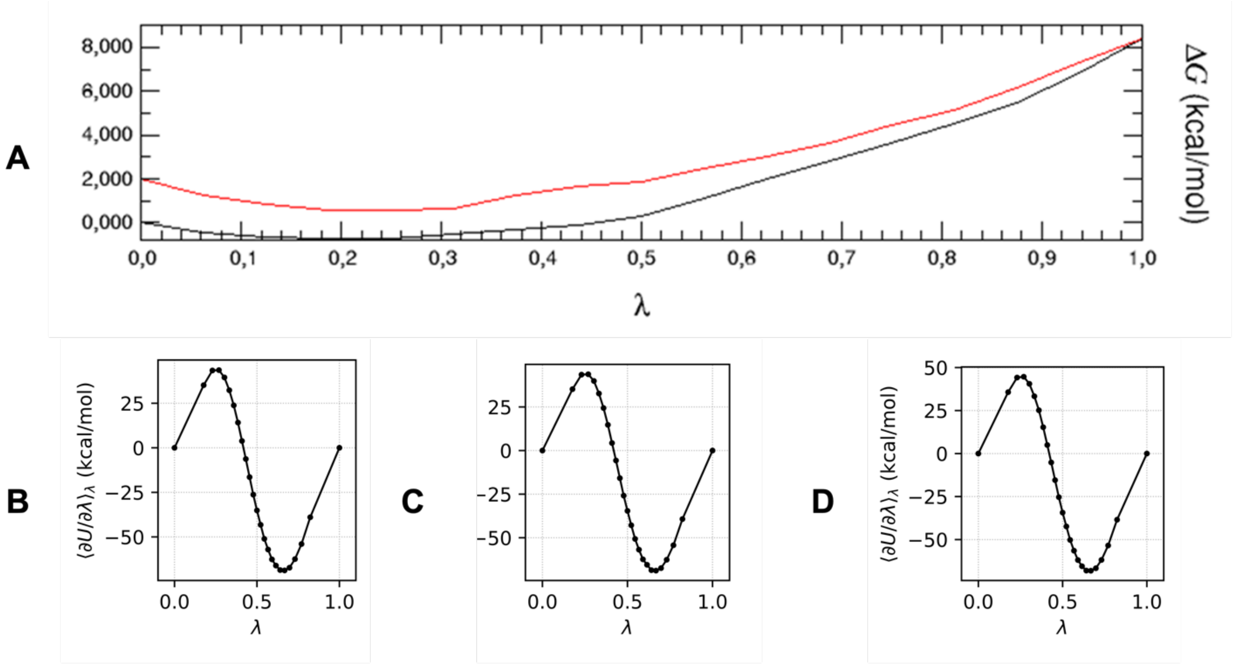


**Figure S21.** A) Free energy (ΔG) versus the coupling parameter λ for the forward (black solid line) and backward (red solid line) perturbation of the perturbation of intermediate-2 to glutamate in the complex phase obtained from NAMD2.14/OPLS-AA/FEP protocol. Variation of the gradient of the potential energy (∂U/∂λ) as a function of λ versus the coupling parameter λ for the perturbation of intermediate-2 to glutamate in the complex phase obtained from B) replica1 of the AMBER20/ff14SB/TI protocol, C) replica2 of the AMBER20/ff14SB/TI protocol, and D) replica3 of the AMBER20/ff14SB/TI protocol.

(3) Karatzas, E.; Zamora, J. E.; Athanasiadis, E.; Dellis, D.; Cournia, Z.; Spyrou, G. M. ChemBioServer 2.0: An Advanced Web Server for Filtering, Clustering and Networking of

Chemical Compounds Facilitating Both Drug Discovery and Repurposing. *Bioinformatics* **2020**, *36* (8), 2602–2604. https://doi.org/10.1093/bioinformatics/btz976.
